# Targeting Metabolic Dysfunction and Inflammation with Sotagliflozin Reverses Diastolic Dysfunction in Experimental HFpEF

**DOI:** 10.1101/2025.11.27.691021

**Authors:** Parisa Shabani, Tahra Luther, Rajesh Chaudhary, Anand Singh, Afnan Alzamrooni, Olga Grushko, Nathan Levine, Rachel Lopez-Schenk, Sascha N. Goonewardena, Bertram Pitt, Ahmed Abdel-Latif

**Affiliations:** Division of Cardiology, Department of Internal Medicine, Frankel Cardiovascular Center, University of Michigan, Ann Arbor, MI 48105, USA; Ann Arbor VA Healthcare System, 2215 Fuller Rd, Ann Arbor, MI 48105, USA

## Abstract

**Background:** Heart failure with preserved ejection fraction (HFpEF) is increasingly prevalent and strongly associated with cardiometabolic comorbidities including obesity, hypertension, and metabolic dysfunction. While SGLT2 inhibitors have demonstrated clinical benefits in HFpEF, the mechanisms underlying dual SGLT1/2 inhibition remain incompletely understood.

**Methods:** We utilized a murine model of cardiometabolic HFpEF induced by high-fat diet combined with L-NAME administration. Following disease establishment, mice received sotagliflozin (30 mg/kg) or vehicle for 10 weeks. Comprehensive assessments included echocardiography, indirect calorimetry, cardiac metabolomics, bulk RNA sequencing with cell-type deconvolution, and high-dimensional immune profiling by flow cytometry and CyTOF.

**Results:** Sotagliflozin significantly attenuated weight gain and improved glucose tolerance without normalizing blood pressure. Metabolic cage analyses revealed a sustained reduction in respiratory exchange ratio, indicating enhanced fatty acid oxidation, corroborated by elevated cardiac acylcarnitine intermediates including palmitoylcarnitine and dodecanoylcarnitine. Echocardiography demonstrated that sotagliflozin protected against diastolic dysfunction, normalizing isovolumic relaxation time and E/e’ ratio while reducing left ventricular mass and myocardial fibrosis. Transcriptomic profiling revealed upregulation of mitochondrial fatty acid β-oxidation pathways and suppression of inflammatory signaling cascades including IL-1 processing and TLR pathways. Flow cytometric analysis demonstrated reduced cardiac infiltration of neutrophils, CCR2+ inflammatory monocytes/macrophages, and IL-1β-expressing immune cells. Splenic immune cell expansion characteristic of systemic inflammation was similarly attenuated.

**Conclusions:** Dual SGLT1/2 inhibition with sotagliflozin exerts coordinated cardiometabolic benefits in experimental HFpEF through metabolic reprogramming toward enhanced lipid utilization and suppression of cardiac and systemic inflammation. These findings establish that sotagliflozin targets the intertwined metabolic-inflammatory axis central to HFpEF pathogenesis, providing mechanistic insight into the therapeutic efficacy of dual SGLT inhibition in cardiometabolic heart failure.

**Graphical abstract:** 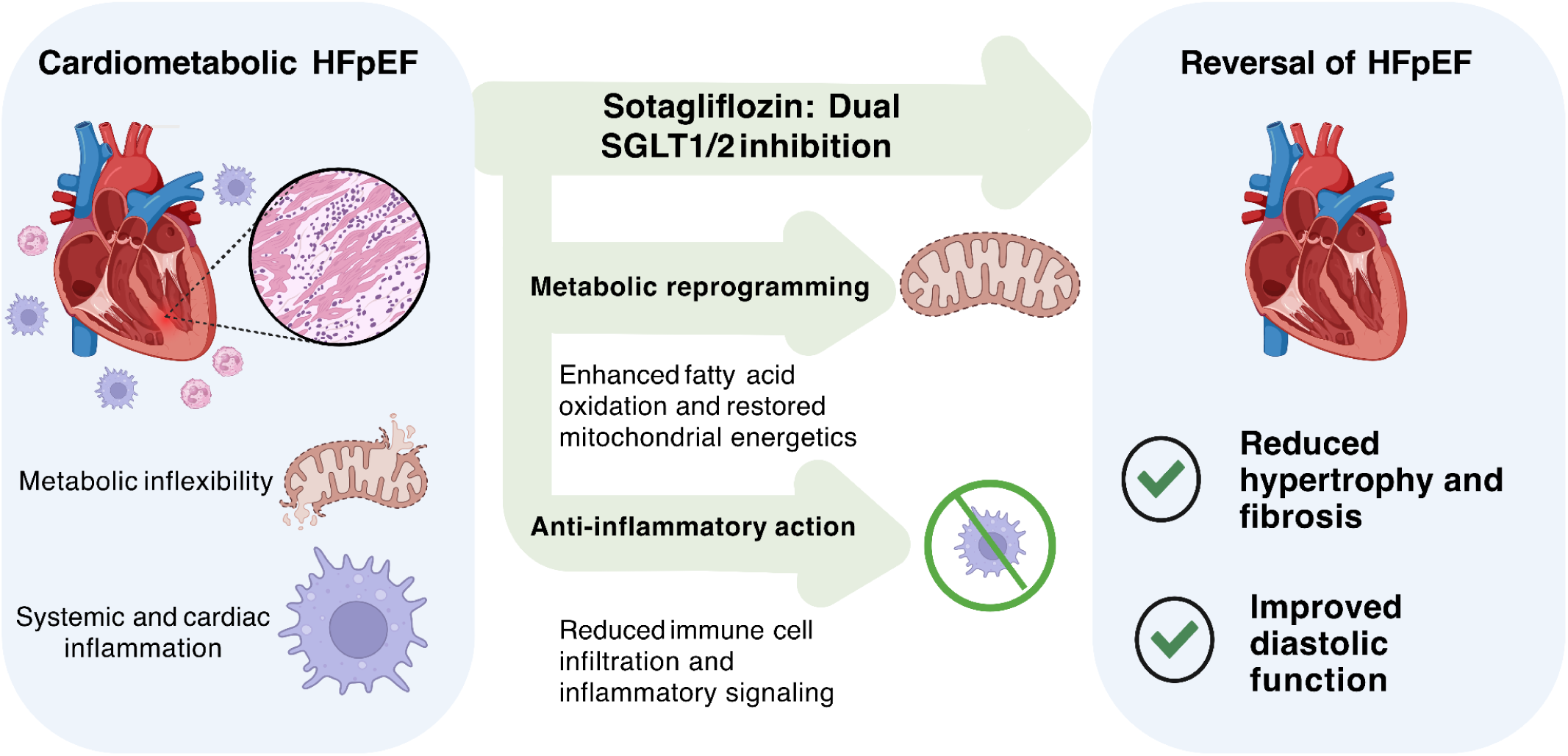

## Introduction

Heart failure (HF) represents a growing global health crisis, affecting millions worldwide and is associated with substantial morbidity, mortality, and healthcare expenditure (Verma et al. 10/2024; Shahim et al. 2023). Approximately 50% of heart failure patients, particularly older adults and women, present with heart failure with preserved ejection fraction (HFpEF), a classification of heart failure characterized by preserved or normal left ventricular ejection fraction (LVEF ≥50%), and diastolic dysfunction or elevated filling pressure (Heidenreich et al. 2022; Dunlay, Roger, and Redfield 2017). Unlike HF with reduced ejection fraction (HFrEF), the prevalence of HFpEF is increasing and carries a similarly poor prognosis; however, therapeutic options that provide mortality benefits have remained elusive until recently (Shah et al. 2017; Anker et al. 2021; Fragasso et al. 2025; Verma et al. 10/2024).

The pathophysiology of HFpEF is complex and heterogeneous. Still, it is strongly linked to the presence of multiple cardiometabolic comorbidities, including hypertension, obesity, type 2 diabetes mellitus (T2DM), and chronic kidney disease (Borlaug 2014; Borlaug and Redfield 2011). These conditions converge to promote systemic inflammation, endothelial dysfunction, coronary microvascular rarefaction, and myocardial metabolic derangement (Franssen et al. 2016). This pro-inflammatory and metabolically stressed environment contributes directly to the key pathological features of HFpEF, such as cardiomyocyte hypertrophy, increased myocardial stiffness due to interstitial fibrosis, impaired relaxation (diastolic dysfunction), and ultimately, elevated left ventricular filling pressures leading to HF symptoms (Paulus and Tschöpe 2013). Consequently, therapeutic strategies targeting the underlying mechanisms of inflammation and metabolic dysfunction are required for the management of HFpEF.

Sodium-glucose cotransporter 2 (SGLT2) inhibitors, initially developed as glucose-lowering agents for T2DM, have emerged as a transformative therapy in cardiovascular medicine (Zelniker and Braunwald 2020). Large-scale clinical trials have demonstrated their robust efficacy in reducing HF hospitalizations and cardiovascular mortality in patients with T2DM at high cardiovascular risk (Zinman, Lachin, and Inzucchi 2016; Neal et al. 2017), and subsequently in patients with HFrEF, irrespective of diabetes status. Recently, landmark trials such as EMPEROR-Preserved and DELIVER have established the benefits of SGLT2 inhibitors in reducing HF hospitalizations or cardiovascular death in patients with HFpEF (Anker et al. 2021; Solomon et al. 2022). The mechanisms underlying these profound cardiovascular benefits are multifactorial and remain incompletely elucidated. Proposed mechanisms include inducing glycosuria and natriuresis, leading to modest reductions in blood pressure and plasma volume, promoting a shift towards ketone body utilization as an alternative cardiac fuel source, improving endothelial function, and exerting direct anti-inflammatory and anti-fibrotic effects within the myocardium (Lytvyn et al. 2017; Packer 2023).

Sotagliflozin is distinct among SGLT inhibitors, as it inhibits SGLT2 and SGLT1 (Powell et al. 2015, 2020; Posch et al. 2022). While SGLT2 is predominantly expressed in the proximal tubules of the kidney and accounts for ∼90% of renal glucose reabsorption, SGLT1 is highly expressed in the small intestine, where it mediates dietary glucose and galactose absorption, and also contributes to the remaining ∼10% of renal glucose reabsorption (Hummel et al. 2012; Powell et al. 2020). Inhibition of intestinal SGLT1 by Sotagliflozin reduces postprandial glucose excursions and may offer additional metabolic benefits beyond SGLT2 inhibition alone. Recent evidence implicates myocardial SGLT1 as an active driver of cardiac injury in diabetic and non-diabetic settings, where its upregulation promotes oxidative stress, sodium overload, fibrosis, and adverse remodeling. Although the degree to which contemporary SGLT2 inhibitors engage SGLT1 remains unknown, convergent mechanistic and genetic data highlight myocardial SGLT1 as a promising therapeutic target in HF and ischemic heart disease (Sayour et al. 2021). Clinical trials like SOLOIST-WHF and SCORED demonstrated that Sotagliflozin reduced cardiovascular death and HF events in patients with diabetes, including those recently hospitalized for worsening HF or with chronic kidney disease (A. S. Bhatt et al. 2024; D. L. Bhatt et al. 2024, 2021). However, the specific contributions of combined SGLT1/2 inhibition to cardiac structure, function, metabolism, and inflammation in the context of cardiometabolic disease, a key driver of HFpEF, remain to be elucidated.

Given the frequent co-existence of hypertension and metabolic dysfunction in HFpEF, we utilized a murine model combining a high-fat diet (HFD) to induce obesity and metabolic derangement with chronic administration of the nitric oxide synthase inhibitor L-NAME to cause hypertension. This model recapitulates key features of human HFpEF, including cardiac hypertrophy, diastolic dysfunction, preserved ejection fraction, and systemic inflammation (Schiattarella et al. 2019). We hypothesized that treatment with the dual SGLT1/2 inhibitor Sotagliflozin, initiated after the disease phenotype was established, would ameliorate cardiac hypertrophy and diastolic dysfunction in this HFD/L-NAME model of hypertensive HFpEF. We further postulated that these benefits would be associated with improvements in glucose metabolism, a shift in systemic substrate utilization, and attenuation of cardiac and systemic inflammation. To test this hypothesis, we performed comprehensive assessments of metabolic parameters, systemic blood pressure, cardiac structure and function, and cardiac and splenic inflammatory markers to delineate the effects of dual SGLT1/2 inhibition in this relevant preclinical model of cardiometabolic HFpEF.

## Methods

### Animals

All animal experiments were performed in accordance with the approved University of Michigan Institutional Animal Care and Use Committee (IACUC) protocol. Sex-specific differences in responses to cardiometabolic syndrome are known to exist in both mice and humans (Salinero, Anderson, and Zuloaga 2018; Goyal et al. 2017). Since female mice are less susceptible to developing cardiac phenotype than males (Tong et al. 2019), we used 8-10-week-old male C57BL/6 mice for this study (Jackson Laboratory, Bar Harbor, ME). Mice underwent a two-week acclimatization period, and baseline measurements of body weight and echocardiogram were taken before they were randomized to receive a low-fat diet (LFD, #D12450J, Research Diet, USA) or a high-fat diet (HFD, #D12492, Research Diet, USA) supplemented with nitric oxide synthase inhibitor, Nω-nitro-L-arginine methyl ester (L-NAME, 0.5 g/L or 100 mg/200 ml) (#N5751, Sigma, USA) in drinking water. After confirming the phenotype at 5 weeks, animals were randomized to receive Sotagliflozin (SOTA) or vehicle at 30 mg/kg once daily via oral gavage. All mice were housed at an ambient temperature of 23 °C, at a density of 2-3 mice/cage in 12 h of light:12 h of darkness with lights off at 07:00 (Zeitgeber, ZT 0) and lights on at ZT 12, and had ad libitum access to food and water throughout the 15-week study duration. Food, water, and cages were changed twice weekly for all mice.

### Intraperitoneal glucose tolerance test (IPGTT)

An intraperitoneal glucose tolerance test (IPGTT) was performed at weeks 5 and 15 after 5 hours of fasting, using 2 g/kg body weight (3-6 mice/group). Blood glucose readings were measured at 0, 15, 30, 60, 90, and 120 minutes after the bolus dose via tail-vein prick using Accu-Chek® Performa II (Roche, USA). The area under the curve (AUC) was calculated using the trapezoidal rule (Allison et al. 1995).

### Body weight and composition

Body weight was recorded weekly during cage, food, and water changes. Body composition was measured in a subset of mice (n=3/group) using an NMR-based analyzer, EchoMRITM-500 Body Composition Analyzer, according to the manufacturer’s instructions, at the University of Michigan Phenotyping Core.

### Sable Metabolic cage studies

At week 14, a subset of HFD + L-NAME mice (n=5/group) was acclimatized for a week to single housing conditions before transferring to Promethion (Comprehensive, High-Resolution Behavioral Analysis Systems, Sable Systems International, USA), at the University of Michigan Animal Phenotyping Core. Food and water consumption, x, y, and z beam breaks, VO2, and VCO2 were measured at 20-minute intervals. Energy expenditure (EE) and respiratory quotient (RQ) were calculated using the Weir equation (Weir 1949). EE was adjusted to raw EE/ (body weight) as previously described (Tschöp et al. 2011).

### Blood pressure measurements

Blood pressure measurements were conducted using a CODA Monitor (Kent Scientific) via the tail-cuff method, according to the manufacturer’s instructions, after the mice were acclimatized and settled in a quiet procedure room with no disturbance. Briefly, an occlusion (O) tail cuff and a volume pressure recording (VPR) cuff were placed on the tail of the mouse, and the O-cuff was inflated to impede blood flow to the tail before the cuff was deflated slowly, and the return blood flow was measured by using a VPR sensor. Body temperature was measured at the base of the tail using a laser thermometer. All measurements were recorded in non-anesthetized mice, and body temperature was maintained using a thermal pad.

### Echocardiography measurements

All echocardiographic measurements were recorded in anesthetized mice maintained with 1-2% isoflurane and 95% oxygen to maintain the heart rate at 450±50 beats per minute. Briefly, trans-thoracic echocardiography was conducted using the Vevo F2 Imaging System (#53699-20, FUJIFILM VisualSonics, Inc., USA). The anterior chest hair was removed using Nair Hair-removal cream before sedation. Body-temperature-maintained, prewarmed ultrasound gel was applied to the area underlying the heart after the mice were immobilized on the stage. The parasternal short-axis view, identified by the presence of the papillary muscles, was used to obtain M-mode images for ejection fraction, fractional shortening, left ventricular mass, cardiac output, and other cardiac parameters. The apical four-chamber view was used to get the tissue Doppler and mitral valve pulse-wave Doppler measurements for myocardial tissue and blood flow velocity, respectively. All parameters were measured at least three times, and average measurements were plotted using their means for statistical analysis.

### Histopathology

Histopathology services were performed by the In Vivo Animal Core Histology Laboratory within the Unit for Laboratory Animal Medicine at the University of Michigan. Briefly, tissues were fixed in 4% paraformaldehyde (PFA) for 24 hr, then transferred to 70% ethanol. Fixed tissues were embedded in paraffin and sectioned at 4 μm thickness on a rotary microtome (TissueTek VIP5®, Sakura Finetek USA, Inc., Torrance, CA).

Following deparaffinization and hydration with xylene and graded alcohols, formalin-fixed, paraffin-embedded (FFPE) slides were placed in 60 °C Bouin’s Fluid (Rowley Biochemical; #F-367-1), a mordant, for 1 hour. Slides were briefly cooled and then rinsed well with water. Slides were stained in Biebrich Scarlet-Acid Fuchsin (Rowley Biochemical; #F-367-3) for 10 minutes, then briefly rinsed in deionized water before being placed in Phosphomolybdic-Phosphotungstic Acid Solution (Rowley Biochemical; #F-367-4), a mordant, for 15 minutes. Finally, slides were transferred directly into Aniline Blue Solution (Rowley Biochemical; #F-367-5) for 8 minutes, followed by a brief rinse in deionized water. Slides were dehydrated and cleared through graded alcohols and xylene, then cover-slipped with Micromount (Leica Biosystems; #3801731) using a Leica CV5030 automatic cover slipper.

### Flow Cytometry and CyTOF Maxpar

Phenotypic cell analysis of macrophages, T cells, and neutrophils in the heart was performed. Briefly, mice were sacrificed, and hearts were rapidly isolated and placed in ice-cold PBS (VWR International). Using a razor blade, the heart was minced manually. After mincing, the tissue was incubated with a mixture of Collagenase type I and type XI (Sigma) and DNase I (Sigma) at 37°C for 30 minutes, with gentle agitation on a rocker. Enzymatic digestion was quenched with cold staining buffer, and the suspension was placed on ice, filtered through a 70 μm cell strainer, then centrifuged at 400 x g for 5 mins at 4°C, followed by debris removal and red blood cell (RBC) lysis steps. The supernatant was discarded, the pellet was resuspended in 1 ml of staining buffer, and the cells were counted. Approximately one million cells were aliquoted per sample for staining.

Cells for flow cytometry were incubated immediately with surface-stain conjugated primary antibodies against RB780conjugated Ly6G (BD Pharmingen), BUV563-conjugated F4/80 (BD), AF488-conjugated LYVE1 (eBioscience), BUV615-conjugated CD11b (BD), APC-CY7-conjugated CD45 (Biolegend), PE-Cy5-conjugated CD64 (Biolegend), BV711-conjugated CD68 (Biolegend), BUV661-conjugated mCCR2 (BD), BV421-conjugated CD115 (Biolegend), BV605-conjugated CD161 (Biolegend), BV570-conjugated CD3 (Biolegend), BUV805-conjugated CD8a (Cytek Biosciences), BUV737-conjugated CD4 (eBioscience) and BUV496-conjugated CD19 (Cytek Biosciences) for 30 minutes on ice. Cells were then washed with staining buffer, permeabilized, and fixed at room temperature for 15 minutes, and incubated with an intracellular marker stain mix of conjugated primary antibodies against PE-eF610-conjugated CD206 (eBioscience), eF450-conjugated IL-1b (eBioscience), and BV650-conjugated TNFa for 20 minutes on ice. Cytek Aurora Spectral Analyzer then analyzed samples in the University of Michigan Flow Cytometry Core. Using unstained cells and single fluorescent controls, laser calibration and compensation were performed for all experiments. CD45hi/Ly6Glo/F4-80hi cells were identified as macrophages and further classified as pro-inflammatory or reparative based on CD206 and CD11c expression. Neutrophils were defined as CD45hi/CD115lo/Ly6-C/G-lo.

Cells for CyTOF staining were counted, and 1-3million cells were aliquoted per sample and barcoded via the Cell-ID 20-Plex Pd Barcoding Kit (Standard Biotools, cat# 201060) to label each sample with their own unique identifying barcode number. Barcoded samples were combined and incubated with a Maxpar metal-conjugated antibody cocktail of Nd-conjugated TCRB, CD115, CD69, F4/80, CD11b, and CD25, Sm-conjugated CD19, CD3e, and TER119, Eu-conjugated CD64, Er-conjugated CD8a and NK1.1, Tm-conjugated CD206, Yb-conjugated CD86, Ly-6G, and CD45, Bi-conjugated CD11c, Gd-conjugated CD62L, Ho-conjugated LYVE1, Lu-conjugated CCR2, and Gd-conjugated CD4 for 30 minutes at room temperature. The concatenated sample was then fixed and permeabilized with Maxpar Fix and Maxpar Perm buffers, respectively, and stained with intracellular Pr-conjugated TNFa and Tb-conjugated IL-1b for 30 minutes at room temperature, then fixed again with 3.2% PFA solution for 10 minutes at room temperature, and intercalated with IrDNA Intercalation and Permeabilization solution for up to overnight at 4 °C. The sample was then washed and resuspended in Maxpar Acquisition solution and run on a Helios mass cytometer from Standard BioTools using CyTOF Software version 7.1.16401.

### Metabolomics

Whole hearts were immediately flash-frozen for metabolomic analysis. The tissue was prepared as previously described (Nwosu et al. 2020). Briefly, the cardiac tissue was lysed in 80% ice-cold methanol using a TissueLyser II (Qiagen) set to 28 Hz, applied in 30-second intervals until a homogeneous mixture was obtained. The samples were pelleted at 13,000 rpm for 5 minutes, and the supernatant was processed in a SpeedVac to get a dry pellet. The dry was reconstituted in 50% methanol and analyzed by LC-MS as detailed in Kerk et al. (Kerk et al., 2022). LC-MS analysis was performed using an Agilent 1290 Infinity II LC –6470 Triple Quadrupole (QqQ) tandem mass spectrometer (MS/MS) system. Data were median-centered and normalized to the sham average. Fold change was calculated by using normalized data.

### RNAseq analyses

Total mRNA was extracted from 20-30 mg of cardiac tissue using TRI Reagent (T9424, Sigma Aldrich, St. Louis, MO, USA) and TissueLyser LT (Qiagen, Hilden, Germany) at 50 Hz. The concentration and purity of mRNA were assessed using a NanoDrop 2000 Spectrophotometer (Thermo Scientific, USA) before synthesizing cDNA from 1000 ng of total RNA from each sample using the Maxima First Strand cDNA Synthesis Kit (#K1671, Thermo Scientific) as per the kit’s protocol on the T100 Thermal Cycler (Applied Biosystems).

### RNA Sequencing and Differential Gene Expression Analysis

Total RNA was extracted from the heart using the RNeasy Plus Mini Kit (Qiagen), following the manufacturer’s instructions. Novogene performed RNA library preparation and transcriptome sequencing. After quality control, 150 bp paired-end sequencing was performed using the Illumina PE150 platform. Transcriptomic data analysis and visualization were performed using Pluto (https://pluto.bio). Paired-end FASTQ files were processed with the nf-core/rnaseq pipeline (v3.17.0). Briefly, reads were trimmed with Trim Galore, aligned to the Mus musculus GRCm39 reference genome using STAR, and quantified to gene-level counts with RSEM.

Differential gene expression analysis was conducted using the DESeq2 R package, which applies a negative binomial model. Genes with fewer than 3 counts in at least 20% of samples in any group were filtered out before analysis. Log2 fold changes were calculated. Statistical significance was determined using a false discovery rate (FDR)-adjusted p-value threshold of 0.05. Volcano plots display log2 fold changes on the x-axis and -log10 adjusted p-values on the y-axis. Genes with an adjusted p-value ≤ 0.05 and a fold change < –0.6 or > 0.6 were considered significantly altered. Up to 12 of these genes were highlighted on the volcano plot.

### Gene Set Enrichment Analysis (GSEA)

Gene set enrichment analysis was performed using the fgsea R package and the fgseaMultilevel() function. Genes were ranked by log2 fold change from the DESeq2 output. C2: Canonical pathways - REACTOME gene set collection from the Molecular Signatures Database (MSigDB) was curated using the msigdbr R package. Gene sets were filtered to include only those containing 5-1,000 genes. Normalized enrichment scores (NES) represent the magnitude and direction of enrichment: positive NES values (shown in red) indicate enrichment in the first group, whereas negative NES values (shown in blue) indicate enrichment in the second group. Statistical significance was assessed using p-values.

### Cell Deconvolution of Bulk RNA-seq Data Using BayesPrism

To perform cellular deconvolution of bulk RNA-seq data, we applied the BayesPrism R package (v2.2.2) in accordance with the authors’ instructions and package vignette (https://github.com/Danko-Lab/BayesPrism). As a reference single-cell dataset, we used publicly available single-cell RNA-seq data from the Gene Expression Omnibus (GSE249412), generated from the same mouse model of HFpEF. Cell types were annotated using canonical markers of major cardiac populations. Because further subdivision into subpopulations was not required for this study, we used identical labels for “cell state” and “cell type.” Lowly expressed genes were removed using the default BayesPrism outlier filtering parameters (outlier.cut = 0.01, outlier.fraction = 0.1). The processed single-cell reference and bulk matrices were then supplied to BayesPrism through the new.prism() function with default settings, and deconvolution was performed using the run.prism() function. Reconstructed expression profiles were subsequently obtained with the get.exp() function.

### In Vitro Assessment of the Anti-Inflammatory Effects of Sotagliflozin in Bone Marrow–Derived Macrophages

We evaluated the anti-inflammatory effects of Sotagliflozin using bone marrow–derived macrophages (BMDMs) isolated from mice. Following differentiation, BMDMs were pre-incubated with varying concentrations of Sotagliflozin for 1 h, after which lipopolysaccharide (LPS, 50 nM) was added for 4 h at 37 °C in a 5% CO₂incubator. Culture supernatants were collected to quantify IL-6 levels using an ELISA kit (BD Biosciences, San Diego, CA, USA) according to the manufacturer’s instructions.

### Statistical analysis

Data are presented as mean ± SEM. The data were checked for normality using the Shapiro-Wilk test, and all statistical analyses were performed on log-transformed data if not normally distributed using GraphPad Prism (Version 10.2.2 (341)). Two-way or repeated measures ANOVA with Bonferroni’s post hoc analysis was performed for group comparison. P<0.05 was considered statistically significant. Metabolomics raw data were processed using the Agilent software (MassHunter Qual and ProFinder). Data analysis was performed with the Agilent MassProfiler Pro package using a recursive analysis workflow.

## Results

### Sotagliflozin Attenuates Metabolic Dysfunction, Weight Gain, and Hypertension in a Murine Model of Hypertensive Heart Failure

To mimic key aspects of human hypertensive heart failure, particularly heart failure with preserved ejection fraction (HFpEF), often associated with metabolic comorbidities, we established a model combining a high-fat diet (HFD) with chronic L-NAME administration to induce hypertension (**Figure 1A**). Consistent with the metabolic challenge, mice receiving the HFD/L-NAME regimen exhibited significantly accelerated weight gain compared with those maintained on a standard low-fat diet (LFD) over the initial 5 weeks and throughout the 15-week study (**Figure 1B**). Therapeutic intervention with the dual SGLT1/2 inhibitor Sotagliflozin (Sotagliflozin), initiated after the establishment of obesity and hypertension (week 5), effectively curbed further weight gain, specifically in the HFD/L-NAME group, resulting in significantly lower body weight by study end compared to vehicle-treated HFD/L-NAME counterparts. Sotagliflozin had no significant impact on the weight trajectory of LFD-fed mice (**Figure 1B**).

**Figure 1.**
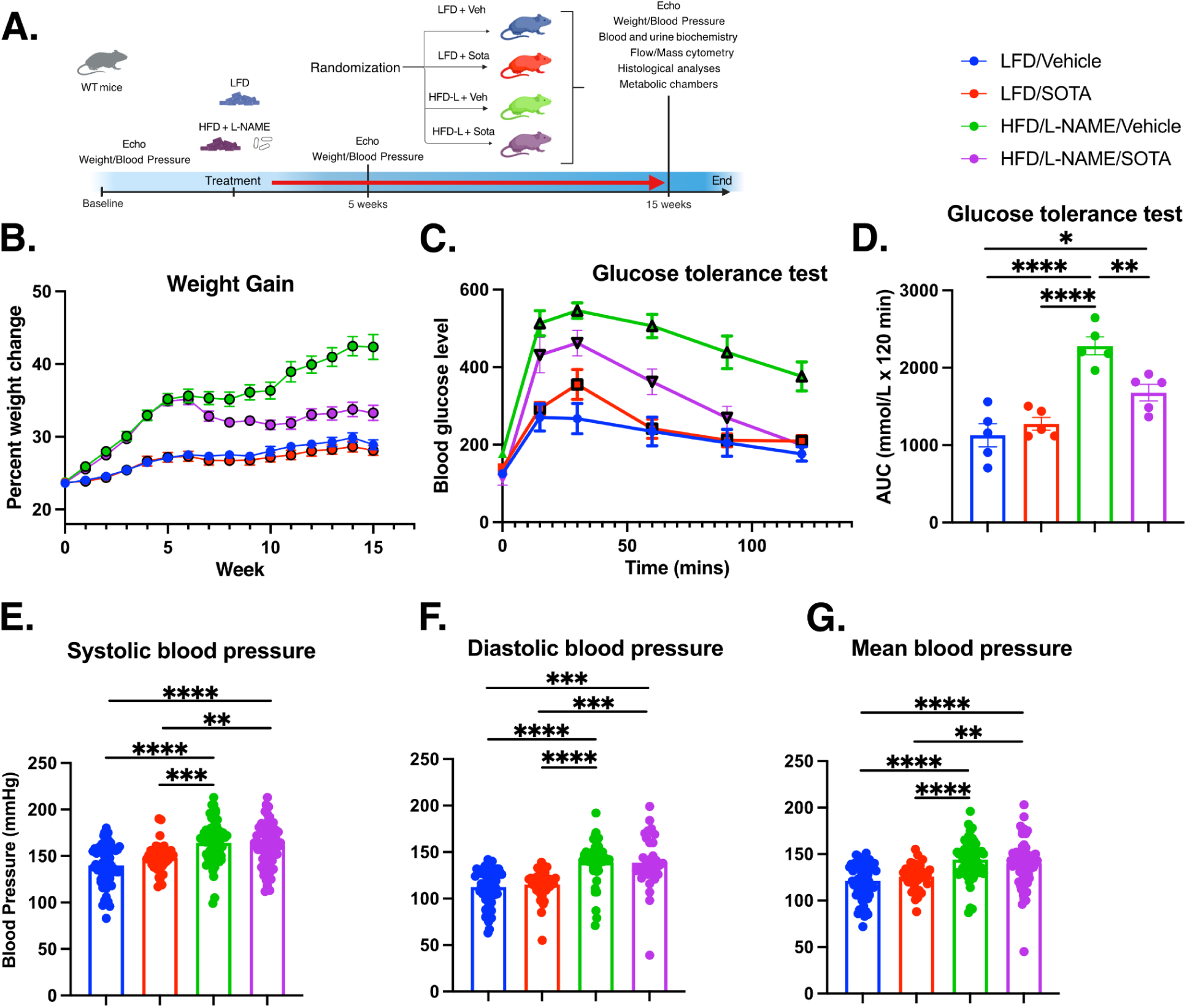
Sotagliflozin attenuates weight gain, improves glucose tolerance, and modestly impacts blood pressure in a diet-induced cardiometabolic heart failure with preserved ejection fraction mouse model. **(A)** Schematic diagram of the experimental protocol. Wild-type (WT) mice were fed either a low-fat diet (LFD) or a high-fat diet combined with L-NAME in drinking water (HFD/L-NAME) for 5 weeks. Baseline echocardiography (Echo), body weight, and blood pressure measurements were taken. Mice were then randomized to receive either vehicle (Veh) or the SGLT1/2 inhibitor Sotagliflozin (SOTA) for 10 weeks, resulting in four groups: LFD/Vehicle, LFD/Sotagliflozin, HFD/L-NAME/Vehicle, and HFD/L-NAME/Sotagliflozin. Endpoint measurements included Echo, weight, blood pressure, blood/urine biochemistry, flow/mass cytometry, histology, and metabolic chamber analyses at 15 weeks. **(B)** Percent weight gain over the 15-week study period for all four experimental groups. **(C)** Blood glucose levels during an intraperitoneal glucose tolerance test (GTT) performed at the study endpoint (week 15). **(D)** Area under the curve (AUC) quantification for the GTT shown in (C). **(E-G)** Endpoint measurements (week 15) of systolic blood pressure (E), diastolic blood pressure (F), and mean blood pressure (G). Data in B-G are presented as mean ± SEM. Statistical significance was determined by one-way or two-way ANOVA, as appropriate, with Bonferroni post hoc tests. *P<0.05, **P<0.01, ***P<0.001, ****P<0.0001. LFD indicates low-fat diet; HFD, high-fat diet; L-NAME, N(G)-nitro-L-arginine methyl ester; Sotagliflozin, Sodium-glucose cotransporter inhibitor (Sotagliflozin); Veh, Vehicle; AUC, Area Under the Curve.

A critical component of the HFpEF phenotype is metabolic dysregulation. We assessed glucose homeostasis via an intraperitoneal glucose tolerance test (GTT) at the study endpoint. The HFD/L-NAME regimen induced severe glucose intolerance, characterized by markedly elevated fasting glucose levels and a pronounced, sustained excursion in blood glucose following the glucose challenge compared to LFD controls (**Figure 1C**). Quantification of the area under the glucose curve (AUC) confirmed this profound impairment in glucose handling in the HFD/L-NAME vehicle group (**Figure 1D**). Treatment with Sotagliflozin dramatically improved glucose metabolism in HFD/L-NAME mice; it lowered fasting glucose and significantly blunted the glycemic response to the glucose load, leading to a considerably reduced AUC, indicative of enhanced glucose disposal and/or reduced glucose absorption (**Figure 1C, 1D**).

As intended by the L-NAME administration, the HFD/L-NAME protocol successfully induced significant hypertension. At week 15, vehicle-treated HFD/L-NAME mice displayed markedly elevated systolic, diastolic, and mean arterial blood pressures compared to normotensive LFD controls, with no significant differences between vehicle and sotagliflozin-treated groups. (**Figure 1E-G**). This result demonstrates that established hypertension in this model was not attenuated by Sotagliflozin, suggesting that blood pressure-independent mechanisms mediate the dual SGLT1/2 inhibitor’s protective effects on metabolic parameters and weight.

### Sotagliflozin Induces a Metabolic Shift Towards Fat Utilization

To understand the metabolic mechanisms underlying Sotagliflozin’s effects, particularly its impact on energy balance and substrate preference, we performed indirect calorimetry using metabolic cages. HFD/L-NAME mice treated with Sotagliflozin exhibited a significantly and consistently lower Respiratory Exchange Ratio (RER = VCO2 produced / VO2 consumed) throughout the 48-hour measurement period compared to vehicle-treated HFD/L-NAME controls (**Figure 2A**). An RER closer to 0.7 indicates preferential oxidation of fats, whereas an RER closer to 1.0 indicates carbohydrate oxidation. The observed sustained reduction in RER strongly suggests that Sotagliflozin promoted a metabolic shift, increasing the reliance on fatty acids as an energy source. This interpretation was directly supported by calculations of substrate oxidation rates, which revealed significantly higher fat oxidation rates (Figure 2E) and correspondingly lower glucose oxidation rates(**Figure 2D**) in the Sotagliflozin-treated HFD/L-NAME group. This shift is consistent with the known mechanism by which SGLT inhibitors induce glycosuria, leading to calorie loss and compensatory mobilization and utilization of fat stores. Despite the apparent shift in substrate use and the observed reduction in body weight gain (**Figure 1B**), overall, 24-hour energy expenditure was not significantly different between the Sotagliflozin and vehicle-treated HFD/L-NAME groups (**Figure 2B**). Body composition analysis showed trends towards reduced fat mass in Sotagliflozin-treated mice, aligning with the overall weight difference (**Figure 2C**). As expected from the osmotic diuresis associated with SGLT inhibition, Sotagliflozin treatment was also associated with a slight increase in water intake (**Figure 2F**). Thus, Sotagliflozin effectively reprogrammed systemic metabolism to favor lipid oxidation, a key mechanism contributing to its beneficial effects in this HFpEF model.

**Figure 2.**
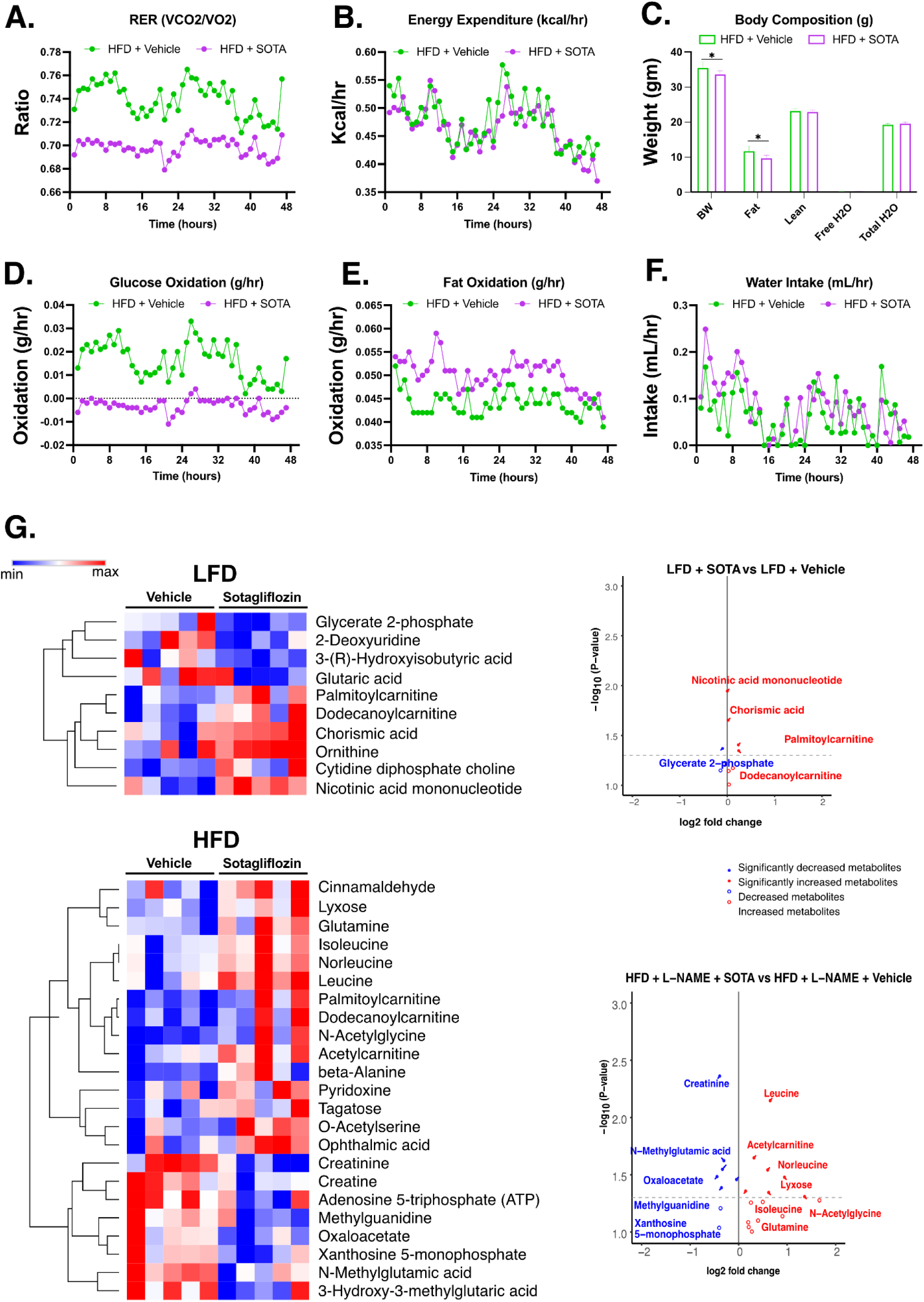
Sotagliflozin shifts substrate metabolism towards fat utilization in HFD/L-NAME mice. Metabolic cage analysis was performed at the study endpoint (week 15) to compare HFD/L-NAME mice treated with vehicle (HFD) versus Sotagliflozin (HFD +). **(A)** Respiratory exchange ratio (RER = VCO2/VO2) over 48 hours. *Note: Y-axis units are dimensionless ratios.* **(B)** Energy expenditure (kcal/hr) over 48 hours. **(C)** Body composition analysis showing total body weight (BW), fat mass, lean mass, free water (H2O), and total body water (H2O) in grams **(D)** Calculated glucose oxidation rate (g/hr) over 48 hours. **(E)** Calculated fat oxidation rate (g/hr) over 48 hours. **(F)** Water intake (mL/hr) over 48 hours. (**G**) Hierarchical clustering heatmap showing metabolites significantly altered (p<0.1) by sotagliflozin treatment in cardiac tissue from low-fat diet (LFD, left panel) and high-fat diet (HFD, right panel) mice. Left panel displays 10 metabolites altered in LFD sotagliflozin-treated (LFDSG) versus vehicle-treated (LFDV) groups. The right panel displays 23 metabolites altered in HFD sotagliflozin-treated (HFDSG) versus vehicle-treated (HFDV) groups. Color scale represents row-normalized metabolite abundance (blue = lower, red = higher). Both panels show distinct metabolic responses to sotagliflozin treatment, with HFD conditions exhibiting more extensive metabolic remodeling, including alterations in fatty acid oxidation (palmitoylcarnitine, dodecanoylcarnitine, acetylcarnitine), BCAA metabolism (leucine, isoleucine, norleucine), energy metabolism (ATP, creatinine, creatine), and TCA cycle intermediates (oxaloacetate, N-methylglutamic acid). LFD conditions show selective changes, including enhanced fatty acid oxidation, NAD+ biosynthesis (nicotinic acid mononucleotide), and membrane remodeling (CDP-choline) with preserved energy homeostasis. Data in A, B, D, E, F are shown as time-course plots (mean values at each time point). Data in C are presented as mean ± SEM. Statistical significance for C was determined using an unpaired t-test, \**P* < 0.05. HFD indicates a high-fat diet with L-NAME/Vehicle group; HFD + SOTA indicates a high-fat diet with L-NAME/Sotagliflozin group.

Our metabolomic studies on cardiac tissue corroborated the metabolic chamber results. In high-fat diet (HFD) mice, sotagliflozin treatment significantly altered 23 cardiac metabolites (p<0.1) compared to vehicle-treated controls, with 65% upregulated (**Figure 2G**). Sotagliflozin treatment enhanced fatty acid β-oxidation relative to vehicle, with palmitoylcarnitine increased 3.73-fold (p=0.053), dodecanoylcarnitine increased 2.22-fold (p=0.073), and acetylcarnitine increased 1.25-fold (p=0.023). Concurrent with enhanced fatty acid metabolism, carbohydrate metabolism was altered in sotagliflozin-treated hearts, with lyxose increased 1.78-fold (p=0.033) and tagatose increased 1.33-fold (p=0.079) compared with vehicle, suggesting increased flux through the pentose phosphate pathway and alternative carbohydrate-processing routes. Branched-chain amino acid (BCAA) metabolism was also activated in sotagliflozin-treated animals, with leucine (1.65-fold, p=0.007), isoleucine (1.63-fold, p=0.045), and norleucine (1.65-fold, p=0.029) all elevated relative to vehicle. Energy-related metabolites showed significant changes: ATP was reduced to 0.78-fold (p=0.092) and xanthosine 5-monophosphate was decreased to 0.78-fold (p=0.033) in sotagliflozin-treated hearts compared with vehicle. Creatine phosphate metabolism was also altered, with creatinine reduced to 0.75-fold (p=0.004) and creatine to 0.98-fold (p=0.035) in the treatment group. TCA cycle intermediates were affected, including reduced oxaloacetate (0.81-fold, p=0.028) and N-methylglutamic acid (0.82-fold, p=0.024), while glutamine increased (1.10-fold, p=0.044) in sotagliflozin-treated animals (**Figure 2G**).

In low-fat diet (LFD) mice, sotagliflozin treatment altered 10 cardiac metabolites (p<0.1) compared to vehicle controls, with 60% upregulated. Sotagliflozin treatment enhanced fatty acid β-oxidation relative to vehicle, with palmitoylcarnitine increased 1.89-fold and dodecanoylcarnitine increased 1.74-fold. Unlike HFD animals, acetylcarnitine was not significantly altered in sotagliflozin-treated LFD mice. Glucose metabolism was also affected by sotagliflozin treatment, with glycerate 2-phosphate decreased to 0.61-fold (p=0.043) in treated hearts compared to vehicle, indicating suppression of glycolysis. BCAA metabolism remained unchanged in sotagliflozin-treated LFD animals compared to vehicle, with no significant alterations in leucine, isoleucine, or norleucine. Energy status was preserved in sotagliflozin-treated LFD hearts, with no significant changes in ATP, nucleotide pools, or creatine phosphate metabolism compared to vehicle controls. TCA cycle intermediates also remained stable in sotagliflozin-treated LFD animals. However, sotagliflozin treatment in LFD mice upregulated nicotinic acid mononucleotide (1.31-fold, p=0.011) and CDP-choline (1.25-fold, p=0.067) compared to vehicle, indicating activation of NAD+ biosynthesis and membrane remodeling pathways (**Figure 2G**).

These findings demonstrate that sotagliflozin treatment produces different metabolic effects depending on dietary background. In HFD conditions, sotagliflozin enhanced fatty acid oxidation while increasing pentose phosphate pathway activity and also altered BCAA catabolism, energy metabolism, and TCA cycle intermediates compared with vehicle. In LFD conditions, sotagliflozin treatment enhanced fatty acid oxidation while suppressing glycolysis, and activated NAD+ biosynthesis and membrane remodeling pathways. At the same time, energy homeostasis and TCA cycle metabolism remained stable compared to vehicle controls.

### Sotagliflozin Protects Against Cardiac Hypertrophy and Diastolic Dysfunction

We next investigated the effects of HFD/L-NAME and Sotagliflozin on cardiac structure and function using transthoracic echocardiography. The HFD/L-NAME regimen induced significant cardiac remodeling and dysfunction characteristic of hypertensive heart disease and HFpEF. Specifically, vehicle-treated HFD/L-NAME mice developed pronounced diastolic dysfunction, evidenced by a significantly prolonged isovolumic relaxation time (IVRT), reflecting slowed myocardial relaxation (**Figure 3B**), and significantly increased E/A and E/e’ ratios, indicating altered diastolic filling patterns and elevated left ventricular filling pressures, respectively (**Figure 3C, 3D**), compared to LFD controls. Remarkably, treatment with Sotagliflozin significantly attenuated these markers of diastolic dysfunction, normalizing IVRT and considerably reducing both E/A and E/e’ ratios in HFD/L-NAME mice (**Figure 3B-D**).

**Figure 3.**
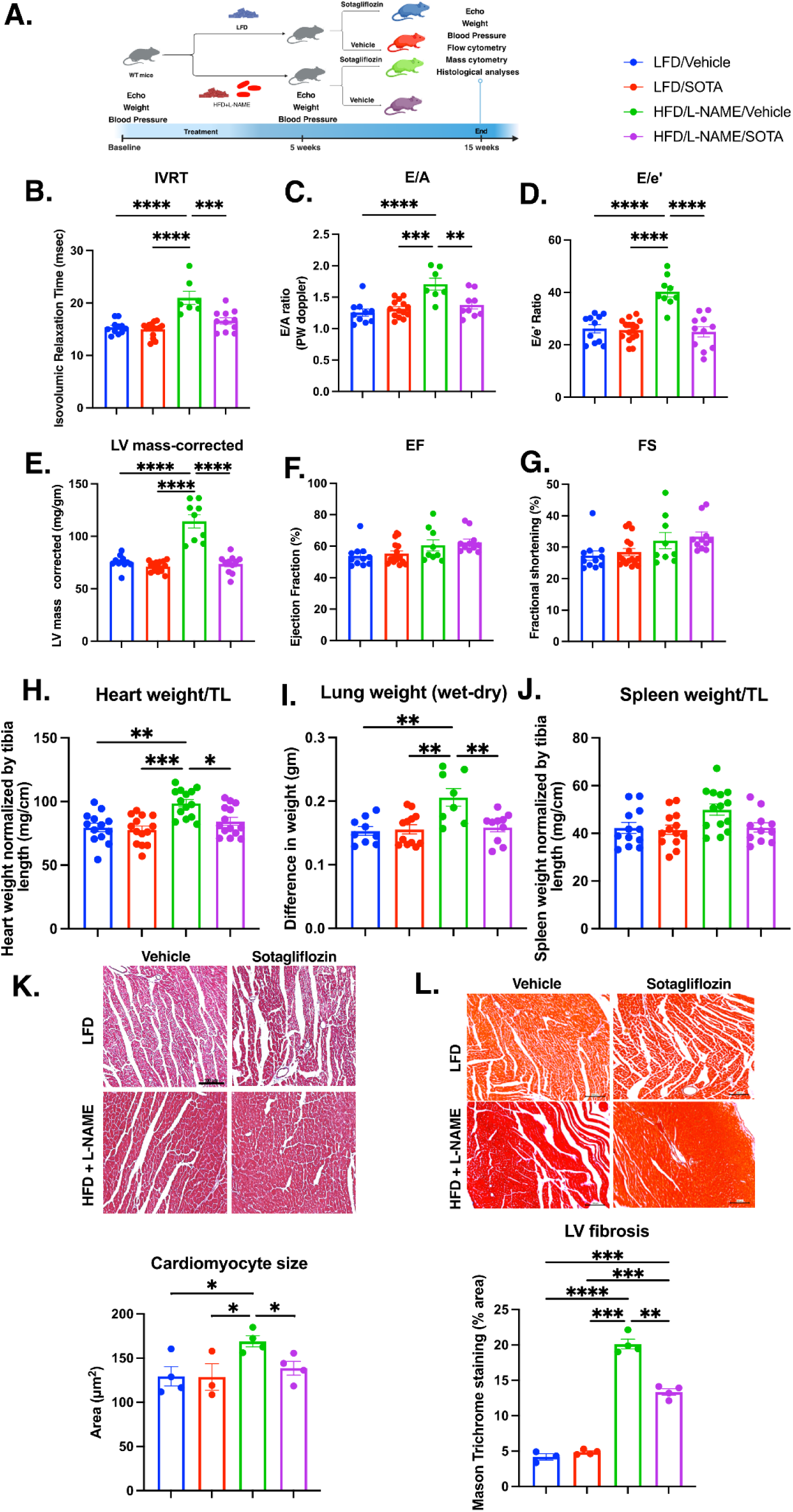
Sotagliflozin ameliorates cardiac hypertrophy, diastolic dysfunction, and pulmonary congestion in HFD/L-NAME mice. Echocardiographic and organ weight analyses at the study endpoint (week 15) across all four experimental groups. **(A)** Schematic diagram of the experimental protocol. **(B)** Isovolumic relaxation time (IVRT, msec). **(C)** Ratio of peak early (E) to late (A) diastolic mitral inflow velocity (E/A ratio), measured by pulsed-wave Doppler. **(D)** Ratio of peak early diastolic mitral inflow velocity (E) to peak early diastolic mitral annular velocity (e’) (E/e’ ratio). **(E)** Left ventricular (LV) mass corrected for body weight (mg/gm). *Note: Correction factor confirmation needed (body weight assumed).* **(F)** LV ejection fraction (EF, %). **(G)** LV fractional shortening (FS, %). **(H)** Gravimetric heart weight normalized to tibia length (TL, mg/cm). **(I)** Lung water content, assessed by the difference between wet and dry lung weight (gm). **(J)** Gravimetric spleen weight normalized to tibia length (TL, mg/cm). (**K**) Assessment of cardiomyocyte size showed a significant reduction in cardiomyocyte surface area with sotagliflozin treatment to a level comparable to that of low-fat diet-treated animals. (**L**) Assessment of left ventricular fibrosis using Mason trichrome staining shows a significant reduction in the total area of cardiac fibrosis in sotagliflozin-treated animals compared to vehicle-treated animals with cardiometabolic stress. Data are presented as mean ± SEM. Statistical significance was determined by two-way ANOVA with Bonferroni post-hoc tests. *P<0.05, **P<0.01, ***P<0.001, ****P<0.0001. LFD indicates low-fat diet; HFD, high-fat diet with L-NAME; Sotagliflozin, Sotagliflozin; Veh, Vehicle; IVRT, Isovolumic Relaxation Time; E/A, Early-to-late diastolic filling ratio; E/e’, Early diastolic filling velocity to mitral annular velocity ratio; LV, Left Ventricular; EF, Ejection Fraction; FS, Fractional Shortening; TL, Tibia Length.

Concomitant with diastolic dysfunction, HFD/L-NAME mice developed significant cardiac hypertrophy in response to the combined pressure overload and metabolic stress. This was demonstrated by a significant increase in left ventricular (LV) mass corrected for body weight (**Figure 3E)** and a considerable increase in total heart weight normalized to tibia length (a measure independent of body weight fluctuations) (**Figure 3H**), compared to LFD controls. Moreover, cardiomyocyte size was increased in the HFpEF group, whereas treatment with sotagliflozin significantly reduced cardiomyocyte size in HFpEF mice (**Figure 3K).** Sotagliflozin treatment provided significant protection against cardiac hypertrophy, markedly reducing both normalized LV mass, normalized heart weight, and cardiomyocyte size in the HFD/L-NAME group (**Figure 3E, 3H, 3K**). Importantly, left ventricular systolic function, assessed by ejection fraction (EF) and fractional shortening (FS), remained preserved mainly across all groups throughout the study (**Figure 3F, 3G**), confirming that the HFD/L-NAME regimen recapitulates a HFpEF-like phenotype rather than HFrEF (heart failure with reduced ejection fraction).

Furthermore, the HFD/L-NAME condition led to increased lung water content (wet-dry weight difference), indicative of pulmonary congestion secondary to cardiac dysfunction. Sotagliflozin treatment significantly reduced this pulmonary congestion (**Figure 3I**). Spleen weight, often reflecting systemic inflammation or extramedullary hematopoiesis, was also increased in HFD/L-NAME mice and was modestly, though significantly, reduced by Sotagliflozin (**Figure 3J**). Importantly, we further observed a significant reduction in left ventricular fibrosis within the Sotagliflozin-treated cohort (**Figure 3L**), indicating a direct impact on adverse cardiac remodeling beyond functional improvements. These findings demonstrate that Sotagliflozin effectively mitigates cardiac hypertrophy, improves diastolic function, and reduces pathological remodeling in this preclinical model of HFpEF.

### SGLT Inhibition Alters the Cardiac Transcriptomic Landscape in a Model of HFpEF

To investigate molecular changes in HFpEF and the impact of SGLT inhibition, we performed transcriptomic analysis of cardiac tissue. Comparison of hearts from the HFpEF vehicle group with those from the LFD vehicle control group revealed extensive alterations in gene expression, indicative of the significant transcriptional reprogramming characteristic of the disease state (**Figure 4A**). Treatment of HFpEF animals with sotagliflozin also induced substantial changes in the cardiac transcriptome compared to vehicle-treated HFpEF animals (**Figure 4B**), suggesting a significant molecular effect of the therapy. Analysis of the overlap between differentially expressed gene (DEG) sets highlighted the scope of these changes (**Figure 4D**). Comparing the HFpEF vehicle to the LFD vehicle (Set C intersection with A and B, plus unique C) identified numerous DEGs associated with the HFpEF phenotype. The comparison of HFpEF Sotagliflozin versus HFpEF Vehicle (Set D and its intersections) identified 750 DEGs directly modulated by Sotagliflozin treatment within the diseased heart. Notably, 375 DEGs were uniquely regulated by Sotagliflozin only in the HFpEF context, representing modulation of genes already altered by the disease state itself (overlapping regions between C and D). Overall, these transcriptomic analyses demonstrate that HFpEF is characterized by profound changes in cardiac gene expression, a signature significantly modulated by Sotagliflozin treatment.

**Figure 4.**
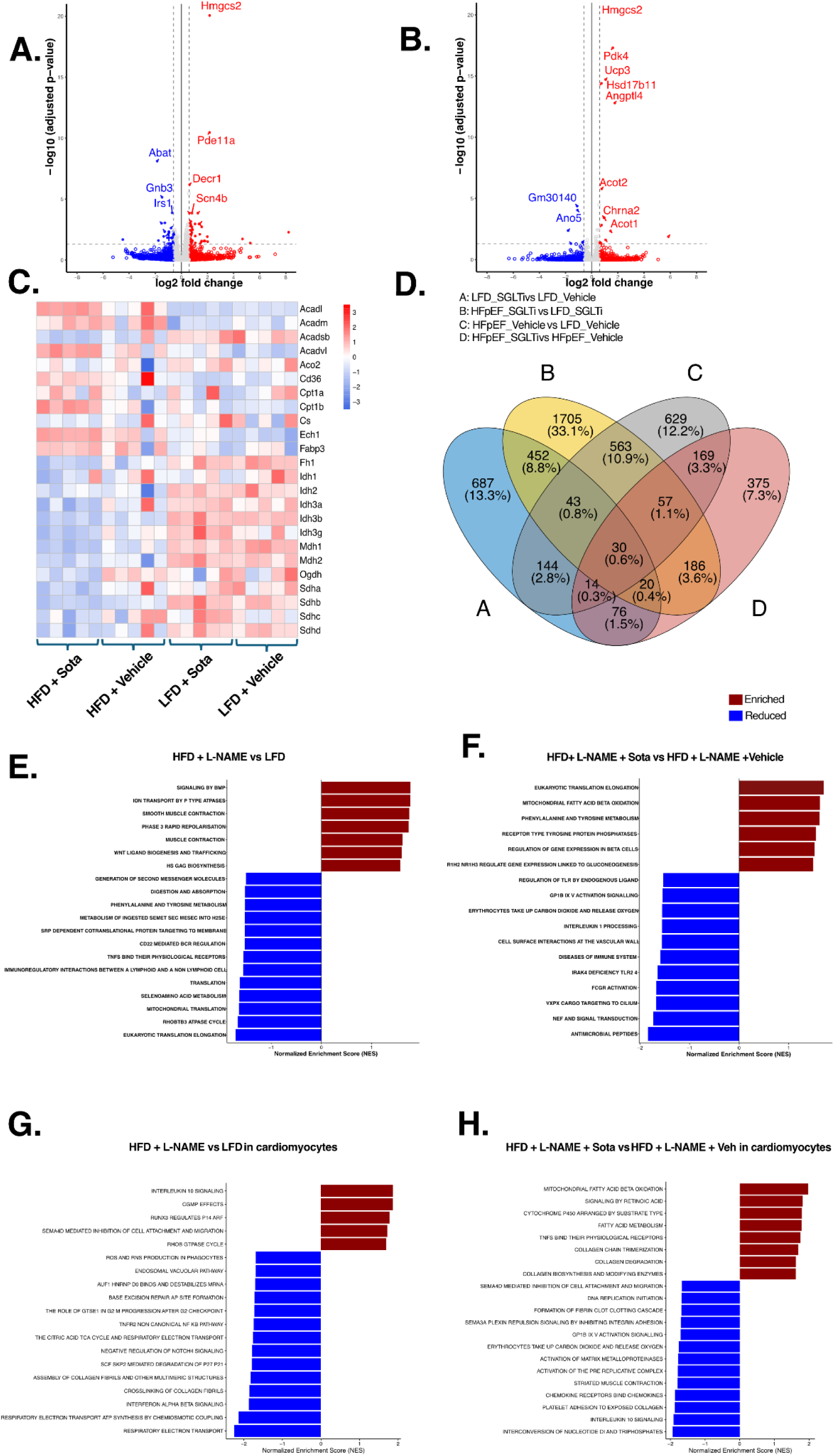
SGLT Inhibition Reverses HFpEF-Induced Cardiac Transcriptomic Alterations, Particularly in Metabolic Pathways. (A, B) Volcano plots showing differentially expressed genes (DEGs) in cardiac tissue comparing (A) HFD + L-NAME vs. LFD vehicle-treated and (B) HFD + L-NAME vs. LFD soatgliflozin-treated. The x-axis represents the log2 fold change, and the y-axis represents the -log10 adjusted p-value. Significantly upregulated genes are shown in red, and downregulated genes are shown in blue (significant threshold: adjusted p-value < 0.05). Key genes are labeled and represent those involved in fatty acid and glucose metabolism and in inflammatory pathways. (C) Heatmap illustrating the expression patterns (Z-score normalized) of selected genes involved in fatty acid, glucose metabolism, and mitochondrial function across four experimental groups: HFD + Sota, HFD + Vehicle, LFD + Sota, and LFD + Vehicle. Red indicates higher relative expression, blue indicates lower relative expression. Overall, HFD + L-NAME was associated with increased fatty acid and glucose metabolism, whereas sotagliflozin reduced glucose metabolism and increased fatty acid metabolism. (D) Venn diagram depicting the overlap of DEGs identified from four comparisons: (A) LFD Sotagliflozin vs. LFD Vehicle, (B) HFpEF Sotagliflozin vs. LFD Sotagliflozin, (C) HFpEF Vehicle vs. LFD Vehicle, and (D) HFpEF Sotagliflozin vs. HFpEF Vehicle. Numbers indicate the count of unique and common DEGs in each intersecting or unique set. (E, F) Gene Set Enrichment Analysis (GSEA) comparing pathway activity in cardiac tissue between (E) HFD + L-NAME vs. LFD and (F) HFD + L-NAME + Sota vs. HFD + L-NAME + Vehicle. Bars represent the Normalized Enrichment Score (NES). Red indicates pathways enriched in the first group of the comparison, while blue indicates pathways depleted (enriched in the second group). The data suggest that HFD + L-NAME is associated with enrichment of pathways involved in muscle hypertrophy, contraction, and ion transport, and reduced activity of amino acid metabolism. Treatment with sotagliflozin enriched pathways involved in gluconeogenesis, glucose metabolism, and amino acid metabolism, with a corresponding reduction in pathways involved in inflammation and immune cell activation. (G, H) GSEA results specifically for cardiomyocyte transcriptomic data, comparing (G) HFD + L-NAME vs. LFD and (H) HFD + L-NAME + Sota vs. HFD + L-NAME + Vehicle. Data suggest increased expression of the inflammatory pathway with HFD + L-NAME, accompanied by induction of the TCA cycle and mitochondrial function pathways. Treatment with sotagliflozin increased fatty acid oxidation and mitochondrial function pathways, while reducing pathways involved in inflammation-related adhesion and chemokine signaling. LFD, low-fat diet; HFD, high-fat diet; HFpEF, heart failure with preserved ejection fraction model (induced by HFD); Sotagliflozin, sodium-glucose cotransporter 2 inhibitor; Vehicle, control treatment; Sota, Sotalol; L-NAME, Nω-nitro-L-arginine methyl ester; DEG, differentially expressed gene; NES, Normalized Enrichment Score.

### HFpEF is Associated with Depressed Cardiac Metabolic Gene Expression, Partially Modulated by SGLT Inhibition

Given the importance of metabolism in cardiac function, we analyzed the expression of key genes involved in fatty acid and mitochondrial metabolism, as shown in the heatmap (**Figure 4C**). Hearts from HFD groups (vehicle and sota, representing the HFpEF condition) generally exhibited reduced expression levels of multiple genes critical for fatty acid beta-oxidation (e.g., *Acadl*, *Acadm*, *Cpt1a*, *Cpt1b*, *Cd36*) and the TCA cycle/mitochondrial respiration (e.g., *Aco2*, *Cs*, *Fh1*, *Idh* family, *Ogdh*, *Sdh* family) compared to LFD controls. In contrast, sotagliflozin in the HFD group led to upregulation of several other genes involved in fatty acid β-oxidation. The overall pattern suggested suppressed oxidative metabolism gene expression in the HFpEF heart relative to controls, along with increased glucose metabolism. This data corroborates the results of the metabolic chamber experiments, identifying a shift in whole body metabolism from glucose utilization to fatty acid oxidation with sotagliflozin treatment. These findings indicate that HFpEF is characterized by a distinctive cardiac metabolic gene expression profile, which is partially restored by Sotagliflozin treatment.

### Pathway Analysis Reveals Broad Metabolic and Signaling Changes in HFpEF and Reversal by SGLT Inhibition

Gene Set Enrichment Analysis (GSEA) was performed to understand the functional consequences of these transcriptomic shifts. In an HFD + L-NAME model, comparison against LFD controls (**Figure 4E**) showed enrichment of some of the signaling pathways (e.g., BMP signaling, ion transport, muscle contraction) but depletion of other pathways related to protein synthesis (Translation, Mitochondrial Translation) in the diseased heart. Sotagliflozin treatment within this model (vs. vehicle, **Figure 4F**) significantly enriched pathways for Mitochondrial Fatty Acid Beta Oxidation and Translation Elongation. Concurrently, Sotagliflozin treatment reduced multiple pathways linked to inflammation and immune activation, including Interleukin-1 processing and TLR signaling.

We then conducted deconvolution analysis of the bulk RNAseq data to explore the contribution of cardiac cells to the overall findings. Focusing specifically on cardiomyocyte transcriptomic data, comparison of HFD + L-NAME versus LFD (**Figure 4G**) revealed enrichment of stress/signaling pathways (e.g., IL-10 signaling, cGMP effects, ROS/RNS production) within the HFpEF cardiomyocytes. Critically, core energy production pathways, including ‘The Citric Acid (TCA) Cycle and Respiratory Electron Transport’ and ‘Respiratory Electron Transport ATP Synthesis’, were markedly reduced in HFpEF cardiomyocytes, confirming the intrinsic mitochondrial dysfunction previously reported. Furthermore, pathways related to collagen assembly were also altered.

Evaluation of sotagliflozin effects specifically within HFpEF cardiomyocytes (HFpEF Sotagliflozin vs HFpEF vehicle, **Figure 4H**) demonstrated a strong enrichment of ‘Mitochondrial Fatty Acid Beta Oxidation’ and overall ‘Fatty Acid Metabolism’ pathways. This indicates that sotagliflozin directly promotes fatty acid utilization pathways within the heart muscle cells themselves in the context of HFpEF. This could be related to substrate availability or improved mitochondrial function. Conversely, Sotagliflozin treatment led to the depletion of several signaling pathways in cardiomyocytes, including those involved in chemokine signaling, platelet adhesion, IL-10 signaling, and DNA replication initiation, suggesting modulation of inflammatory and signaling cascades directly within the cardiomyocyte compartment. In summary, GSEA and deconvolution analyses highlight the significant transcriptomic and functional pathway alterations in HFpEF hearts and cardiomyocytes, with Sotagliflozin treatment effectively modulating key metabolic, inflammatory, and signaling pathways.

### HFpEF Alters the Cardiac Immune Cell Landscape and Activation State

To investigate the role of immune cells in HFpEF and the effects of SGLT inhibition, we characterized cardiac immune cell infiltrates using high-dimensional spectral flow cytometry. Analysis revealed distinct immune cell clusters within the heart (**Figure 5A**), which could be identified based on canonical surface marker expression patterns corresponding to major lineages, including neutrophils, macrophages, B cells, T cells, and NK cells (**Figure 5C**).

**Figure 5.**
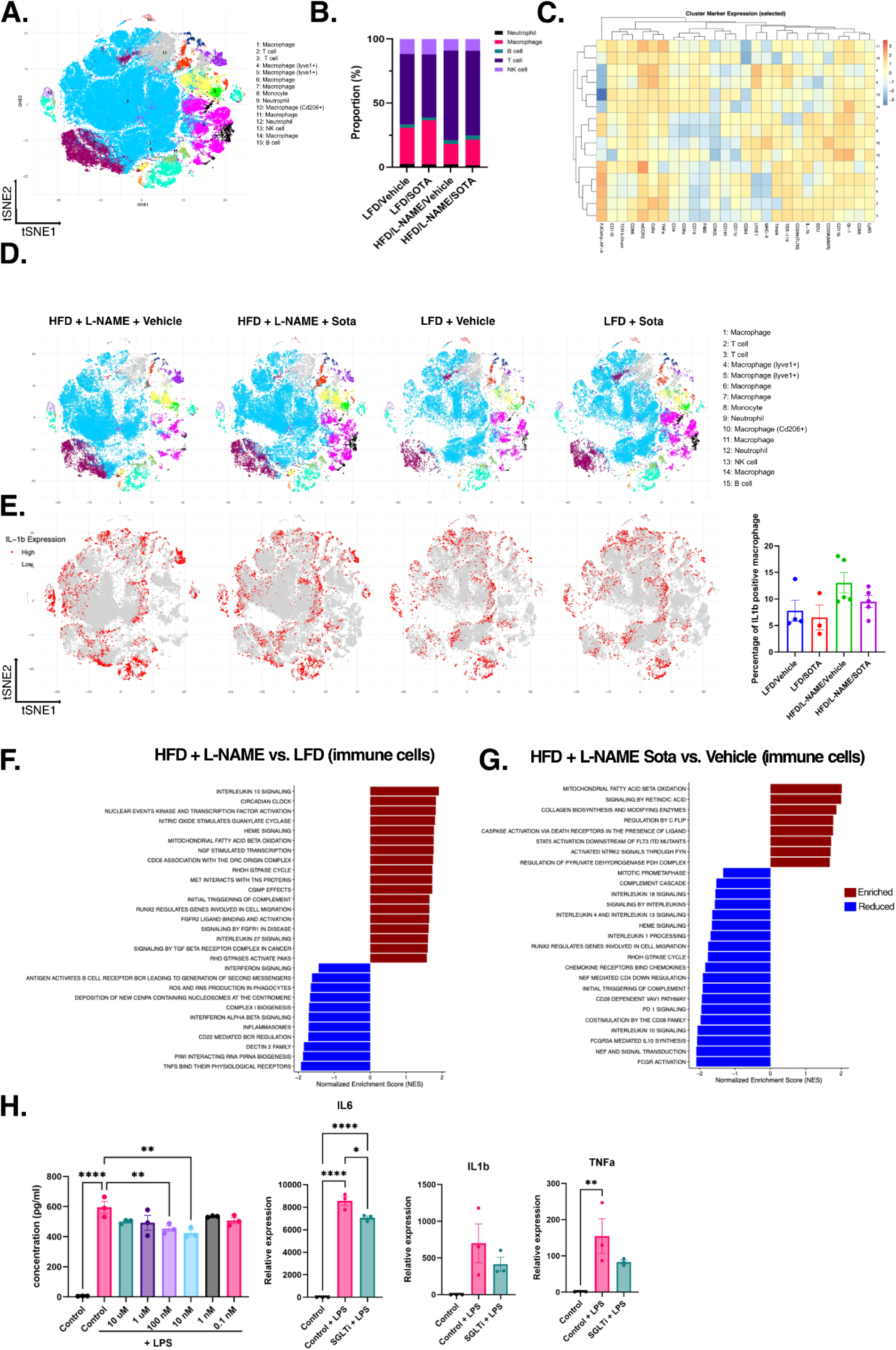
Characterization of Cardiac Immune Cell Infiltrate and Activation States in HFpEF and Modulation by SGLT Inhibition. (A) t-Distributed Stochastic Neighbor Embedding (t-SNE) plot visualizing high-dimensional flow cytometry data of cardiac immune cells. Cells are clustered using FlowSOM, with distinct populations indicated by color. (B) Bar chart showing the average percentage composition of major immune cell types (Neutrophils, Macrophages, B cells, T cells, NK cells) within the total live CD45+ cardiac immune cell gate across experimental groups: LFD Vehicle, HFpEF Vehicle, LFD Sotagliflozin, and HFpEF Sotagliflozin. (C) Heatmap displaying the median expression intensity of selected cell surface markers used for lineage identification and activation state assessment across the immune cell clusters defined in (A). (D) t-SNE plots overlaid with cells from individual experimental conditions (HFD + L-NAME + Vehicle, HFD + L-NAME + Sota, LFD + Vehicle, LFD + Sota), colored by cluster identity as in (A), revealing condition-specific alterations in immune cell population distribution. (E) t-SNE plots corresponding to the conditions in (D), highlighting cells based on IL-1b expression levels (Red = High expression, Grey = Low expression), visualizing the cellular sources and condition-dependent changes in IL-1b production within the cardiac immune infiltrate. This data suggests higher expression of IL-1β among macrophages and T cells. (F, G) Gene Set Enrichment Analysis (GSEA) of pathways altered within the cardiac immune cell populations (derived from inferred pathway activity data), comparing (F) HFpEF Vehicle vs. LFD Vehicle and (G) HFpEF Sotagliflozin vs. HFpEF Vehicle. Bars represent the Normalized Enrichment Score (NES). Red indicates pathways enriched, and blue indicates pathways depleted in the first condition listed in each comparison. SGLT inhibitor treatment exerts significant immunomodulatory effects within the cardiac immune cell population in the HFpEF model. It broadly dampens multiple pro-inflammatory pathways (complement, various interleukins, chemokine signaling) and pathways associated with adaptive immune responses (T-cell co-stimulation, PD-1). Concurrently, it promotes fatty acid metabolism within these cells. This suggests that a key benefit of SGLT inhibition in HFpEF is the reduction of detrimental cardiac inflammation by altering the activation state and metabolic profile of resident and recruited immune cells. LFD, low-fat diet; HFD, high-fat diet; HFpEF, heart failure with preserved ejection fraction model (induced by HFD); Sotagliflozin, sodium-glucose cotransporter 2 inhibitor; Vehicle, control treatment; Sota, Sotalol; L-NAME, Nω-nitro-L-arginine methyl ester; t-SNE, t-Distributed Stochastic Neighbor Embedding; NES, Normalized Enrichment Score; IL-1b, Interleukin-1 beta.

Assessment of the relative proportions of these significant immune populations indicated shifts in the overall cardiac immune cell composition between LFD and HFD + L-NAME groups, with potential further modulation observed following sotagliflozin treatment (**Figure 5B**). Visualizing cells from different experimental conditions on t-SNE maps highlighted distinct distributions, indicating condition-specific alterations in the immune cell landscape (**Figure 5D**, comparing HFD+L-NAME +/- Sota and LFD +/- Sota groups). Notably, examination of Interleukin-1 beta (IL-1b) expression revealed increased populations of IL-1b-expressing cells, particularly evident in the HFD+L-NAME vehicle condition, which were reduced following sotagliflozin treatment (**Figure 5E**), suggesting modulation of inflammatory cytokine production within the cardiac immune infiltrate. In an in vitro study, bone marrow–derived macrophages were treated with varying concentrations of sotagliflozin before LPS stimulation, and IL-6 levels were measured in the conditioned medium. Pre-incubation with sotagliflozin attenuated IL-6 production, with the most potent effects observed at 10 nM and 100 nM. We next evaluated the expression of *Tnfα*, *Il6*, and *Il1b* at the 10 nM dose. IL-6 expression was significantly reduced in the Sotagliflozin + LPS group, while *Tnfα* and *Il1b* exhibited a downward trend compared with LPS alone (**Figure 5H**).

Collectively, these findings demonstrate that HFpEF is associated with significant alterations in the cardiac immune cell landscape, and SGLT inhibition with SOTA modulates this infiltrate, including a reduction in inflammatory cytokine-producing cells.

### Distinct Transcriptional Profiles Characterize Immune Cells in HFpEF

To define the functional state of these immune cells, Gene Set Enrichment Analysis (GSEA) was performed on transcriptomic data derived from cardiac immune populations. Comparison of immune cells from HFD + L-NAME hearts versus LFD controls revealed significant differences (**Figure 5F**). Immune cells in the HFpEF heart showed enrichment in pathways such as ‘Interleukin 10 Signaling’, ‘Heme Signaling’, ‘Mitochondrial Fatty Acid Beta Oxidation’, and ‘NGF Stimulated Transcription’. This indicates a distinct activation signature and potential metabolic reprogramming within the immune cell compartment in the HFpEF setting.

Flow cytometry quantification of digested hearts demonstrated marked remodeling of the cardiac immune compartment in HFD + L-NAME–treated mice, with partial normalization by SGLT inhibition (**Figure 6**). Total CD45⁺ leukocytes were increased in HFD + L-NAME/vehicle hearts relative to LFD/vehicle, consistent with enhanced cardiac inflammation, with a trend towards reduction in HFD + L-NAME/SOTA hearts (**Figure 6B**). This response was driven in part by a robust expansion of neutrophils and Ly6C^hi monocytes in HFD + L-NAME/vehicle hearts (**Figure 6C,E**). Sotagliflozin significantly reduced neutrophil accumulation and tended to lower Ly6C^hi monocytes, while TNFα⁺ neutrophils remained infrequent and unchanged across HFD + L-NAME groups (**Figure 6D**).

**Figure 6.**
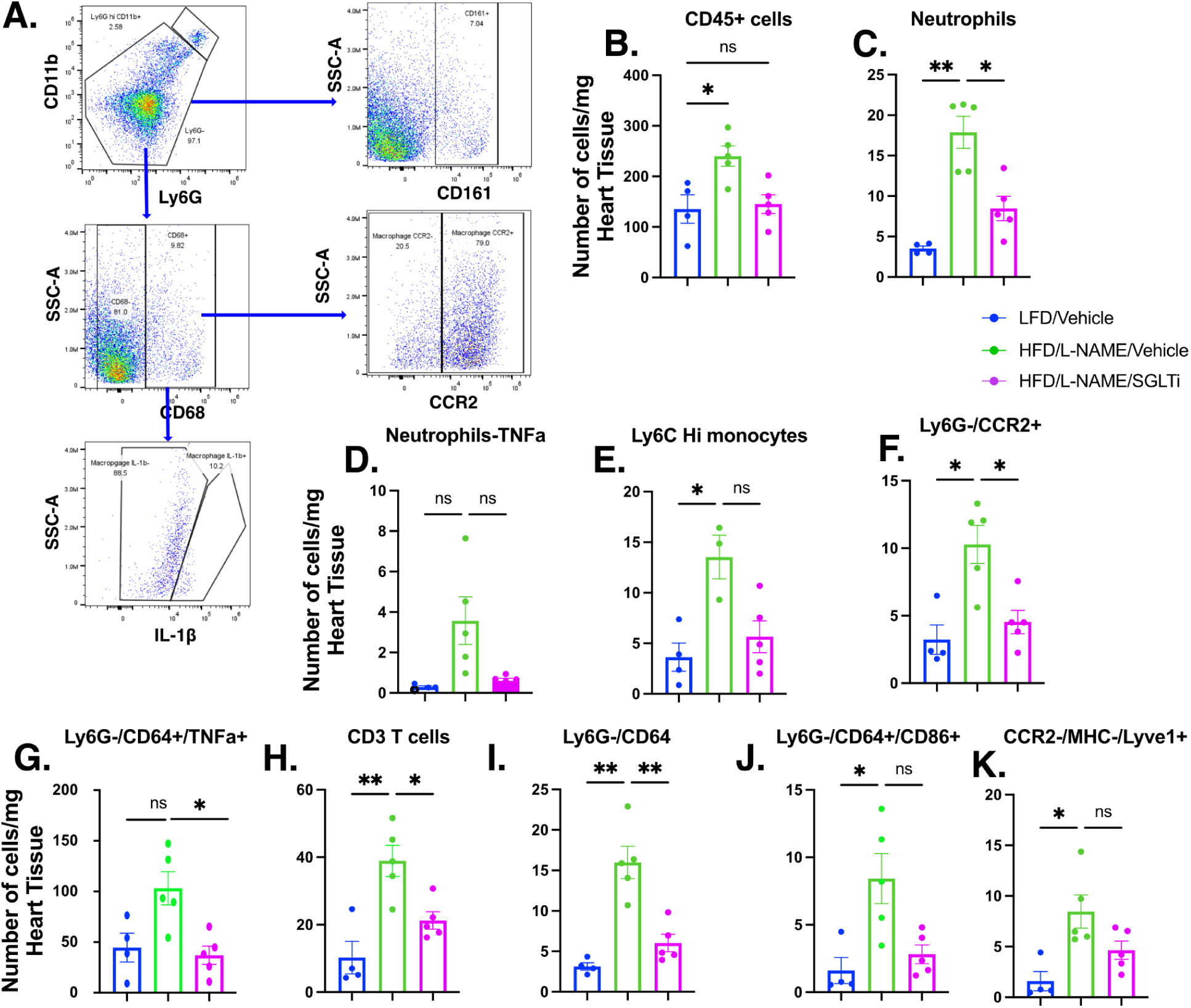
Cardiac immune cell infiltration induced by HFD/L-NAME is differentially modulated by SGLT inhibition. (**A**) Representative flow cytometry gating strategy for cardiac leukocytes. Live singlets were gated on CD45+, followed by identification of CD11b+Ly6G+ neutrophils, Ly6G−CD11b+ myeloid cells, Ly6G−/CD64+ and CD68+ macrophages, CCR2+ inflammatory monocytes/macrophages, IL-1β– or TNFα-producing macrophages, CD3+ T cells, and CCR2−/MHCII−/Lyve1+ resident macrophages, as indicated. (**B–K**) Quantification of leukocyte subsets per mg heart tissue in mice on low-fat diet (LFD)/vehicle, HFD/L-NAME/vehicle, or HFD/L-NAME treated with SGLT inhibitor (HFD/L-NAME/Sotagliflozin). (**B**) Total CD45+ leukocytes are increased in HFD/L-NAME vs LFD, without significant reduction by Sotagliflozin. (**C**) Neutrophil numbers are markedly elevated in HFD/L-NAME hearts and significantly lowered by Sotagliflozin. (**D**) TNFα+ neutrophils are rare and not significantly different among groups. (**E**) Ly6C^hi monocytes and (**F**) Ly6G−/CCR2+ inflammatory monocytes/macrophages are increased in HFD/L-NAME hearts; Sotagliflozin significantly reduces CCR2+ cells but not Ly6C^hi monocytes. (**G**) TNFα+ Ly6G−/CD64+ macrophages show a significant reduction with Sotagliflozin compared with HFD/L-NAME vehicle. (H) CD3+ T cells are expanded in HFD/L-NAME hearts and partially reduced by Sotagliflozin. (**I**) Total Ly6G−/CD64+ macrophages are increased in HFD/L-NAME vs LFD and significantly decreased by Sotagliflozin. (**J**) Pro-inflammatory Ly6G−/CD64+/CD86+ macrophages and (K) CCR2−/MHCII−/Lyve1+ macrophages are elevated in HFD/L-NAME hearts; Sotagliflozin does not significantly alter these subsets. Bars represent mean ± SEM; each symbol denotes an individual mouse. *P<0.05, **P<0.01, ns not significant by one-way ANOVA with Dunnett post hoc multiple comparisons.

HFD + L-NAME/vehicle also increased Ly6G⁻/CCR2⁺ inflammatory monocytes/macrophages (**Figure 6F**) and Ly6G⁻/CD64⁺ macrophages, including CD86⁺ pro-inflammatory macrophages (**Figure 6I,J**), indicating expansion of inflammatory myeloid populations. Sotagliflozin significantly reduced CCR2⁺ cells and total Ly6G⁻/CD64⁺ macrophages, whereas CD86⁺ macrophages showed only a partial, non-significant decrease (**Figure 6F,J**). CD3⁺ T cells were elevated in HFD + L-NAME/vehicle hearts and modestly attenuated with sotagliflozin, while CCR2⁻/MHCII⁻/Lyve1⁺ macrophages were increased by HFD + L-NAME but not significantly altered by sotagliflozin (**Figure 6H,K**). Collectively, these data indicate that HFD + L-NAME provokes broad myeloid and lymphoid infiltration of the heart. SGLT1/2 inhibition preferentially limits neutrophil and CCR2⁺ inflammatory macrophage recruitment, with more modest effects on resident and CD86⁺ macrophage subsets.

### SGLT1/2 Inhibition Modulates Immune Cell Metabolism and Suppresses Inflammatory Pathways in HFpEF

We next evaluated the impact of SGLT inhibition on the immune cell transcriptome within the HFpEF model by comparing HFpEF Sotagliflozin versus HFpEF vehicle groups (**Figure 5G**). Sotagliflozin treatment led to further enrichment of ‘Mitochondrial Fatty Acid Beta Oxidation’ within the immune cells, similar to effects observed in cardiomyocytes, as well as enrichment of ‘Signaling by Retinoic Acid’. Strikingly, SGLT inhibition resulted in the significant reduction of a broad spectrum of pro-inflammatory and immune activation pathways. These included the ‘Complement Cascade’, multiple interleukin signaling pathways (’Signaling by Interleukins’, ‘IL-18 Signaling’, ‘IL-4 and IL-13 Signaling’, ‘IL-1 Processing’), ‘Chemokine Receptors Bind Chemokines’, pathways related to T-cell activation and regulation (’PD-1 Signaling’, ‘Costimulation by the CD28 family’), and pathways linked to phagocytosis and antibody response (’FCGR Activation’). These findings demonstrate that SGLT inhibition significantly dampens inflammatory signaling cascades within cardiac immune cells in the context of HFpEF, while concurrently influencing their metabolic profile. Thus, SGLT inhibition with sotagliflozin profoundly impacts the cardiac immune cell transcriptome in HFpEF, suppressing inflammatory pathways and promoting a metabolic shift towards fatty acid oxidation.

### Sotagliflozin Mitigates Splenic Immune Cell Expansion in HFpEF

To determine if the cardiac effects observed were associated with changes in systemic immune cell distribution, we analyzed immune cell populations in the spleen using Cytometry by Time-Of-Flight (CyTOF) (representative gating strategy shown in Figure 7A). We compared animals on a LFD or a HFD combined with L-NAME to induce HFpEF, treated with either vehicle (veh) or the dual SGLT1/2 inhibitor Sotagliflozin (SOTA).

**Figure 7.**
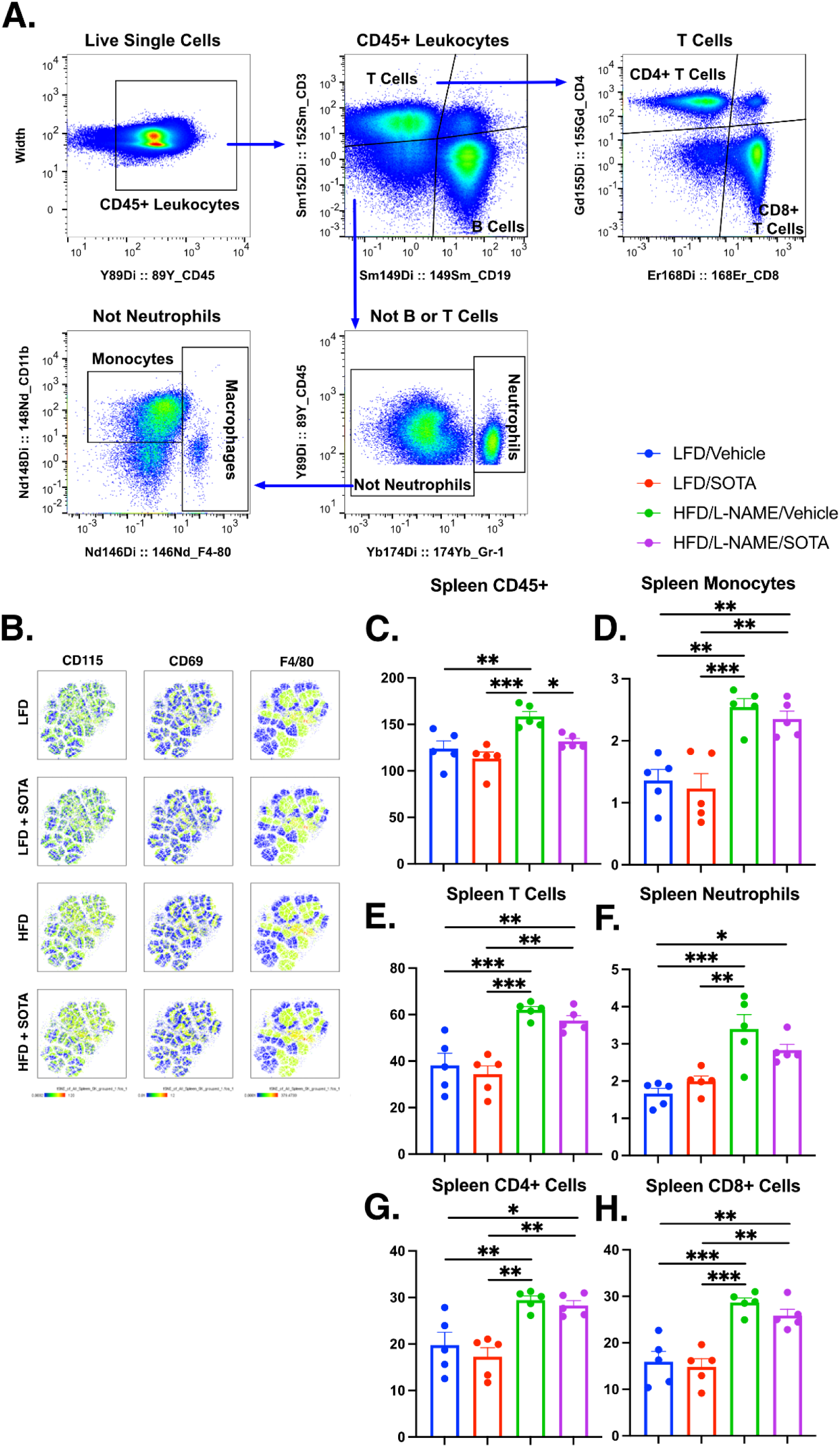
Sotagliflozin Attenuates Systemic Immune Cell Expansion in the Spleen of HFpEF Animals. **(A)** Representative flow cytometry gating strategy for identifying major immune cell populations within live, single splenocytes. Gates define CD45+ leukocytes, T cells (CD3+), B cells (CD19+), CD4+ T cells, CD8+ T cells, monocytes (CD11b+F4/80lo/int), macrophages (CD11b+F4/80hi), and neutrophils (Gr-1hi). **(B)** t-Distributed Stochastic Neighbor Embedding (t-SNE) plots visualizing splenic immune cells from each experimental group (LFD Vehicle, LFD + SOTA, HFD HFD/L-NAME Vehicle, HFD/L-NAME + SOTA). Plots are colored by the expression intensity (Blue=low, Yellow=high) of selected markers: CD115 (CSF1R), CD69 (activation marker), and F4/80 (macrophage marker), highlighting potential shifts in myeloid cell states. **(C-H)** Quantification of the frequency (percentage of parent gate or relative count) of specific immune cell populations in the spleen across the four treatment groups: (C) Total CD45+ leukocytes, (D) Monocytes, (E) Total T cells, (F) Neutrophils, (G) CD4+ T cells, and (H) CD8+ T cells. Data are presented as mean ± SEM. ANOVA determined statistical significance with Bonferroni post-hoc tests: *p < 0.05, **p < 0.01, ***p < 0.001. LFD, low-fat diet; HFD, high-fat diet; L-NAME, Nω-nitro-L-arginine methyl ester; SOTA, Sotagliflozin (SGLT1/2 inhibitor); Veh, Vehicle.

Compared to LFD/vehicle controls, animals in the HFD/L-NAME/vehicle group exhibited significant expansion of multiple splenic immune cell populations, indicative of systemic inflammation. This included a significant increase in the frequency of total CD45+ leukocytes (**Figure 7C**, p<0.001), monocytes (**Figure 7D**, p<0.001), total T cells (**Figure 7E**, p<0.001), neutrophils (**Figure 7F**, p<0.001), CD4+ T cells (**Figure 7G**, p<0.01), and CD8+ T cells (**Figure 7H**, p<0.001). Visual analysis using t-SNE plots suggested accompanying alterations in myeloid cell marker expression profiles, such as F4/80, potentially reflecting changes in macrophage populations or activation states (**Figure 7B**). Treatment with Sotagliflozin significantly attenuated this systemic immune cell expansion in the HFpEF model. Comparing HFD/L-NAME/SOTA animals to HFD/L-NAME/vehicle animals, Sotagliflozin treatment resulted in a significant reduction in the frequency of total CD45+ leukocytes (**Figure 7C**, p<0.05), and a trend towards reduction in other immune cell populations. In contrast, sotagliflozin treatment did not significantly alter the frequencies of these major splenic immune populations in healthy LFD animals (LFD/SOTA vs. LFD/Vehicle, **Figures 7C-H**). These findings indicate that the HFpEF model induces a significant systemic inflammatory response characterized by splenic immune cell expansion, and treatment with the dual SGLT1/2 inhibitor Sotagliflozin modestly mitigates these changes. Overall, these data demonstrate that the HFpEF model is associated with significant systemic immune cell expansion in the spleen, and Sotagliflozin treatment partially attenuates this inflammatory response.

## Discussion

Clinical studies have shown that Sotagliflozin treatment improves clinical outcomes in patients with heart failure with preserved ejection fraction. However, the underlying mechanisms are poorly understood. This study demonstrates that dual SGLT1/2 inhibition with sotagliflozin exerts multifaceted cardiometabolic benefits in a murine model of diet-induced HFpEF. Sotagliflozin therapy, initiated after establishing the HFpEF phenotype, effectively improved glucose homeostasis and modestly reduced blood pressure. Furthermore, Sotagliflozin induced a significant metabolic shift towards fatty acid oxidation at both whole-body and cardiac levels, consistent with its known mechanism of action. Beyond systemic effects, treatment with Sotagliflozin remarkably protected against cardiac hypertrophy and diastolic dysfunction, concurrently mitigating myocardial fibrosis. Mechanistically, these benefits reflect a coordinated reprogramming of energy metabolism toward enhanced lipid utilization, remodeling of amino acid and nucleotide metabolism, and suppression of inflammatory signaling. Together, these effects highlight how dual SGLT inhibition targets the intertwined metabolic and inflammatory pathways central to HFpEF pathogenesis (Anker et al. 2021; Solomon et al. 2022; A. S. Bhatt et al. 2024).

Sotagliflozin-treated HFD/L-NAME mice exhibited marked improvements in diastolic function, evidenced by normalization of isovolumic relaxation time and reductions in E/A and E/e′ ratios, alongside reduced LV mass and myocardial fibrosis. Systolic function remained preserved, confirming a true HFpEF phenotype. The modest decrease in systemic blood pressure was insufficient to account for these cardiac benefits, consistent with clinical findings showing that SGLT2 inhibitors confer disproportionate structural and functional improvements beyond hemodynamic unloading (Anker et al. 2021; A. S. Bhatt et al. 2024; D. L. Bhatt et al. 2024; Pandey et al. 2023). Our results are consistent with previous reports showing improvement in cardiac hypertrophy, fibrosis, and diastolic function following pressure overload using the transaortic constriction (TAC) model (Young et al. 2021). These results emphasize that sotagliflozin’s cardioprotective mechanisms primarily target intrinsic myocardial metabolic and inflammatory processes rather than afterload alone.

Metabolic cage analyses revealed that sotagliflozin induced a sustained reduction in respiratory exchange ratio (RER), indicating a pronounced shift toward fatty acid oxidation. This was supported by significantly higher calculated fat oxidation rates and significant suppression of glucose oxidation, without changes in total energy expenditure. These systemic findings were mirrored in the myocardium, with bulk cardiac tissue transcriptomics and metabolomics demonstrating substantial remodeling of intermediary metabolism consistent with enhanced β-oxidation and altered energy substrate flux. Collectively, these data suggest that sotagliflozin significantly alters systemic and cardiac energy metabolism, favoring fatty acid utilization.

In HFD-fed hearts, sotagliflozin increased multiple acylcarnitine intermediates, including palmitoylcarnitine and dodecanoylcarnitine, indicating accelerated mitochondrial fatty acid import and β-oxidation. This metabolic remodeling was accompanied by modest increases in carbohydrate intermediates such as lyxose and tagatose, suggesting upregulation of the pentose phosphate pathway (PPP), potentially enhancing NADPH production and redox buffering. Consistent with this shift, transcriptomic profiling revealed upregulation of genes associated with mitochondrial fatty acid oxidation (Cpt1b, Acadm, Hadha), reinforcing the metabolic cage results and metabolomics findings (Chen and Su 2015; Szrok-Jurga et al. 2023). Using our deconvolution bioinformatics methods, we traced these transcriptomic changes to cardiomyocytes, highlighting the direct effect of Sotagliflozin on cardiomyocyte metabolism. These data collectively indicate that sotagliflozin enhances mitochondrial lipid utilization and metabolic flexibility, correcting the energy inflexibility characteristic of HFpEF (Stanley, Recchia, and Lopaschuk 2005; Lopaschuk et al. 2021).

Sotagliflozin also modulated amino acid and nucleotide metabolism in HFD-fed animals. Increased levels of branched-chain amino acids (leucine, isoleucine, norleucine) suggest enhanced BCAA catabolism, which could feed anaplerotic substrates into the TCA cycle and support mitochondrial energetics. Meanwhile, reductions in ATP, creatine phosphate, and TCA intermediates such as oxaloacetate may reflect an adaptive downregulation of energy-demand pathways as mitochondrial efficiency improves. The observed increase in glutamine likely supports anaplerotic metabolism and antioxidant defense via glutathione synthesis, which has been shown to enhance cardiomyocyte energetics and function (Alhasan et al. 2024). These findings align with previous research demonstrating that SGLT2 inhibitors redirect metabolic fuel utilization from glucose to lipids (Szekeres, Toth, and Szabados 2021). Together, these findings support a model in which sotagliflozin promotes metabolic reprogramming that reduces glycolytic dependence while enhancing lipid and amino acid–derived energy production, a mechanism consistent with the “energy optimization” paradigm proposed for gliflozins (Mudaliar, Alloju, and Henry 2015; Packer 2023).

In low-fat diet-fed mice, sotagliflozin induced a qualitatively similar but quantitatively muted metabolic response. Fatty acid oxidation markers (palmitoylcarnitine, dodecanoylcarnitine) were increased, while glycolytic intermediates such as glycerate-2-phosphate were suppressed, confirming a systemic shift away from carbohydrate oxidation even under metabolically normal conditions. Notably, NAD⁺-related metabolites, including nicotinic acid mononucleotide and CDP-choline, were upregulated, suggesting stimulation of NAD⁺ biosynthesis and phospholipid remodeling pathways. This observation aligns with prior evidence that SGLT inhibitors enhance NAD⁺ salvage and redox signaling, potentially further improving mitochondrial resilience (Hsu, Seifert, and Arany 2023). Taken together, these comprehensive metabolic data demonstrate that sotagliflozin promotes a profound and beneficial metabolic shift in the heart, enhancing lipid utilization and energy flexibility in HFpEF, a core mechanism likely contributing to its clinical efficacy.

The recent identification of pantothenate kinase 1 (PANK1) as a direct cardiac target of SGLT2 inhibitors (Forelli et al. 2024) provides mechanistic insight into the metabolic reprogramming observed with sotagliflozin in experimental HFpEF. The PANK1-mediated enhancement of coenzyme A biosynthesis elegantly explains our findings of increased cardiac acylcarnitine intermediates (palmitoylcarnitine, dodecanoylcarnitine) and transcriptional upregulation of mitochondrial β-oxidation pathways, suggesting that Sotagliflozin may have similar direct effects to empagliflozin on oxidative fuel metabolism in the failing heart. Our demonstration that sotagliflozin reversed diastolic dysfunction, reduced myocardial fibrosis, and shifted both systemic and cardiac metabolism toward enhanced fatty acid utilization aligns with this cardiomyocyte-intrinsic mechanism, whereby SGLT inhibitor-activated PANK1 restores metabolic flexibility independent of renal or systemic effects.

Collectively, these integrated data indicate that sotagliflozin orchestrates a coordinated metabolic adaptation involving enhanced fatty acid oxidation, activation of the PPP, stimulation of NAD⁺ biosynthesis, and remodeling of amino acid metabolism. These processes collectively improve myocardial energy efficiency, reduce oxidative stress, and restore redox homeostasis in the metabolically stressed HFpEF heart.

In parallel with its metabolic effects, sotagliflozin exerted pronounced anti-inflammatory actions. HFD/L-NAME mice developed systemic inflammation, characterized by splenomegaly and infiltration of monocyte, neutrophil, and T-cell populations into the heart, which was attenuated by sotagliflozin. Transcriptomic profiling of cardiac tissue confirmed broad downregulation of pro-inflammatory pathways, including IL-1β, IL-18, TLR, and complement signaling. These pathways are key effectors of the NLRP3 inflammasome cascade, suggesting that sotagliflozin suppresses inflammasome activation (Byrne et al. 2020; Dai et al. 2023). Given the overlap between metabolic stress and inflammation, the observed metabolic remodeling, particularly increased NAD⁺ and reduced glycolytic flux, may indirectly contribute to anti-inflammatory effects by limiting pro-inflammatory macrophage polarization.

An emerging synthesis links these observations: inflammation itself can induce aberrant cardiac SGLT2 expression, which in turn amplifies oxidative stress and inflammatory signaling (Ali Mroueh et al., 2024; A. Mroueh et al., 2024). By suppressing inflammatory cytokines such as IL-1β and TNF-α, sotagliflozin may interrupt this feed-forward loop, preventing pathological SGLT2 induction and further reducing myocardial oxidative stress. Thus, the anti-inflammatory and metabolic benefits of sotagliflozin are mechanistically interconnected, targeting the same maladaptive axis of metabolic-inflammation crosstalk that defines cardiometabolic HFpEF (Paulus and Tschöpe 2013; Castiglione et al. 2023).

While most SGLT2 inhibitors act predominantly via renal and hemodynamic mechanisms, sotagliflozin’s dual SGLT1/2 blockade confers additional metabolic and potentially direct cardiac effects. Inhibition of intestinal SGLT1 reduces postprandial glucose absorption and insulin excursions (Zambrowicz et al. 2012), indirectly mitigating hyperinsulinemia-driven myocardial hypertrophy (Oktay, Rich, and Shah 2013). Cardiac SGLT1 expression, though debated (Hummel et al. 2012; Pepin, Drakos, and Selzman 2019), may become inducible under inflammatory stress (A. Mroueh et al. 2024; Ali Mroueh et al. 2024), suggesting that dual inhibition prevents maladaptive substrate influx during metabolic overload. Moreover, the observed transcriptional upregulation of fatty acid oxidation genes and elevated acylcarnitines support activation of pantothenate kinase 1 (PANK1) and restoration of coenzyme A pools. This recently described off-target mechanism enhances mitochondrial flux independent of SGLT inhibition (Hsu, Seifert, and Arany 2023). These combined effects, intestinal SGLT1 inhibition, renal SGLT2 blockade, and potential direct myocardial metabolic adaptations to cardiometabolic syndrome, could explain sotagliflozin’s robust metabolic and anti-inflammatory benefits in HFpEF.

## Limitations and Perspectives

The study’s limitations include reliance on the L-NAME model, which induces hypertension by inhibiting NOS and may not fully reflect the features of essential hypertension. Furthermore, HFpEF is a heterogeneous clinical condition with multiple subpopulations. However, the cardiometabolic phenotype of HFpEF, represented in this study, is the fastest-growing subpopulation of this disease. The metabolomic analyses, though comprehensive, were performed in bulk cardiac tissue and cannot distinguish cell-type–specific metabolic shifts. Additionally, the specific contributions of SGLT1 versus SGLT2 inhibition remain to be dissected through selective inhibitor and tissue-specific receptor KO studies in animals. Nevertheless, the consistency between systemic, metabolic, and transcriptional data provides strong mechanistic coherence.

## Conclusions

Dual SGLT1/2 inhibition with sotagliflozin exerts integrated systemic and myocardial benefits in cardiometabolic HFpEF, encompassing metabolic reprogramming, anti-inflammatory activity, and improved diastolic performance. The combination of indirect calorimetry, cardiac metabolomics, and transcriptomics reveals that sotagliflozin enhances fatty acid oxidation, augments NAD⁺-related metabolism, and suppresses inflammasome-mediated signaling. These effects collectively mitigate metabolic inflexibility and inflammation—the two converging drivers of HFpEF pathophysiology. Mechanistically, these findings bridge clinical outcomes with molecular processes, positioning dual SGLT1/2 inhibition as a comprehensive therapeutic strategy for metabolic-inflammatory HFpEF.

## Acknowledgments

Ahmed Abdel-Latif is supported by the VA Merit (I01CX002684-01), NIH R61/R33 (1R61HL177474-01), and the Mathers Foundation grant. Additional funds from Lexicon supported this project.

An AHA postdoctoral fellowship supports Rajesh Chaudhary and Afnan Alzamrooni.

Sascha N. Goonewardena is supported by the Merit Award: 1I01CX002560 (PI) and the NIH award: R01HL150392

## Notes

### Competing Interest Statement

The authors have declared no competing interest.

